# An Expanded Substrate Scope for Cross-Chiral Ligation Enables Efficient Synthesis of Long L-RNAs

**DOI:** 10.1101/2024.10.04.616258

**Authors:** Xuan Han, Jonathan T. Sczepanski

## Abstract

Despite the growing interest in mirror-image L-oligonucleotides, both as a robust nucleic acid analogue and as an artificial genetic polymer, their broader adoption in research and medicine remains hindered by challenges associated with the synthesis of long sequences, especially for L-RNA. Herein, we present a novel strategy for assembling long L-RNAs via the joining of two or more shorter fragments using cross-chiral ligase ribozymes together with new substrate activation chemistry. We show that 5′-monophosphorylated L-RNA, which is readily prepared by solid phase synthesis, can be activated by chemical attachment of a 5′-adenosine monophosphate (AMP) or diphosphate (ADP), yielding 5′-adenosyl (di-or tri-) phosphate L-RNA. The activation reaction is performed in mild aqueous conditions, proceeds efficiently with short or large L-RNA, and, yielding few biproducts, requires little or no further purification after activation. Importantly, both groups, when added to L-RNA, are compatible with ribozyme-mediated ligation, with the 5′-adenosyl triphosphate permitting rapid and efficient joining of multiple, long L-RNA strands. This is exemplified by the assembly of a 129-nt L-RNA molecule via a single cross-chiral ligation event. Overall, by relying on ribozymes that can be readily prepared by *in vitro* transcription and L-RNA substrates that can be activated through simple chemistry, these methods are expected to make long L-RNAs more accessible to a wider range of researchers and facilitate the expansion of L-ON-based technologies.

## INTRODUCTION

L-DNA and L-RNA are the enantiomers of our native genetic polymers, D-DNA and D-RNA, respectively. Although L-oligonucleotides (ONs) do not naturally occur, they can be synthesized chemically in the laboratory, providing researchers with a unique opportunity to build and study mirror-image biology systems. From a biotechnology standpoint, L-ON’s offer several advantages over their native counterparts.^1^ Notably, L-ONs exhibit high resistance to nuclease degradation, which greatly enhances their stability in challenging biological conditions.^2, 3^ L-ONs are also less prone to off-target hybridization with endogenous nucleic acids because Watson-Crick (WC) base pairing is stereoselective.^2, 4, 5^ Moreover, as enantiomers, D- and L-ONs share identical physical and chemical properties, such as thermostability, making them equivalent from a rational design perspective.^4-6^ Due to these beneficial properties, L-ONs are being increasingly utilized in diverse biomedical applications, including molecular imaging^7-10^, clinical diagnostics^11-15^, and aptamer-based therapeutics.^16^

Due to their inverted backbone, L-ONs are not compatible with enzymatic synthesis using native DNA and RNA polymerases.^17^ Consequently, L-ONs are typically prepared using solid-phase phosphoramidite chemistry. This imposes practical limitations on the length and quality of L-ONs that can be currently obtained, especially for L-RNA, placing many of exciting applications mentioned above out of reach. With this in mind, researchers have sought alternative strategies to synthesize long L-ONs. Perhaps the most obvious solution is simply inverting the stereochemistry of natural polymerase enzymes to facilitate L-ON synthesis. Indeed, several groups have successfully prepared D-amino acid versions of protein polymerases^18-23^ and ligases^24^, permitting the assembly and amplification of gene-sized L-DNA fragments and transcription of full-length ribosomal L-RNAs. While this progress is promising, the chemical synthesis of large mirror-image polymerases (>400 amino acids) remains a highly specialized process that is labor intensive, costly, and difficult to scale. Thus, off-the-shelf solutions that are more accessible and practical for the average researcher are still needed.

An alternative approach to using mirror-image enzymes is the use of so called “cross-chiral” enzymes, i.e., enzymes composed of native biopolymers (D-ONs and L-amino acids) that can directly act on L-ONs. Such enzymes could be readily produced using standard biochemical and molecular biology techniques, making them accessible to a wider range of researchers. Towards this goal, Joyce and colleagues have reported a cross-chiral ribozyme, 16-12t, that catalyzes the RNA-templated ligation of RNA molecules of the opposite handedness (Figure 1a).^25^ The ligation reaction involves attack of the 3′-hydroxyl of the “acceptor” RNA substrate on the 5′-triphosphate of the “donor” RNA substrate, resulting in formation of a 3′,5′-phosphodiester linkage. Iterations of this reaction have enabled the assembling long L-RNAs from a mixture of L-trinucleotide building blocks and exponential amplification of L-RNA via “ribo-PCR”.^26, 27^ Thus, cross-chiral assembly of long L-RNAs using D-ribozymes, which are easily obtained by *in vitro* transcription, is well within reach. Nevertheless, this approach also suffers from a key limitation: synthesis of the 5′-triphosphorylated L-RNA donor substrates. The most common method for preparing RNA 5′-triphosphates, *in vitro* transcription^28^, is not possible with enantio-RNA. Thus, 5′-triphosphates must be added to L-RNA through chemical means.^29-35^ Unfortunately, current methods for the chemical triphosphorylation of ONs are often plagued by poor yields and undesired side products. For example, many of these methods use the phosphitylation reagent salicyl phosphorochloridite, which was developed by Ludwig and Eckstein for the solution-phase triphosphorylation of mononucleosides (Figure 1b).^36^ Salicyl phosphorochloridite is highly reactive to water, leading to 5′-H-phosphonate side products if rigorous anhydrous conditions are not maintained.^29, 36-38^ Moreover, this method often results in 10%-20% of the 5′-diphosphate side product due to contamination in the tributylammonium pyrophosphate reagent.^30, 31, 35, 38^ Both of these side products are very difficult to purify from the desired triphosphorylated product, especially for longer oligonucleotides. The yield of the desired 5′-triphosphorylated product also decreases precipitously with increasing ON length.^33, 35^ Taken together, it remains difficult to produce 5′-triphosphorylated L-RNAs in sufficient yield and quality to support the cross-chiral assembly of long L-RNA molecules.

**Figure 1.**
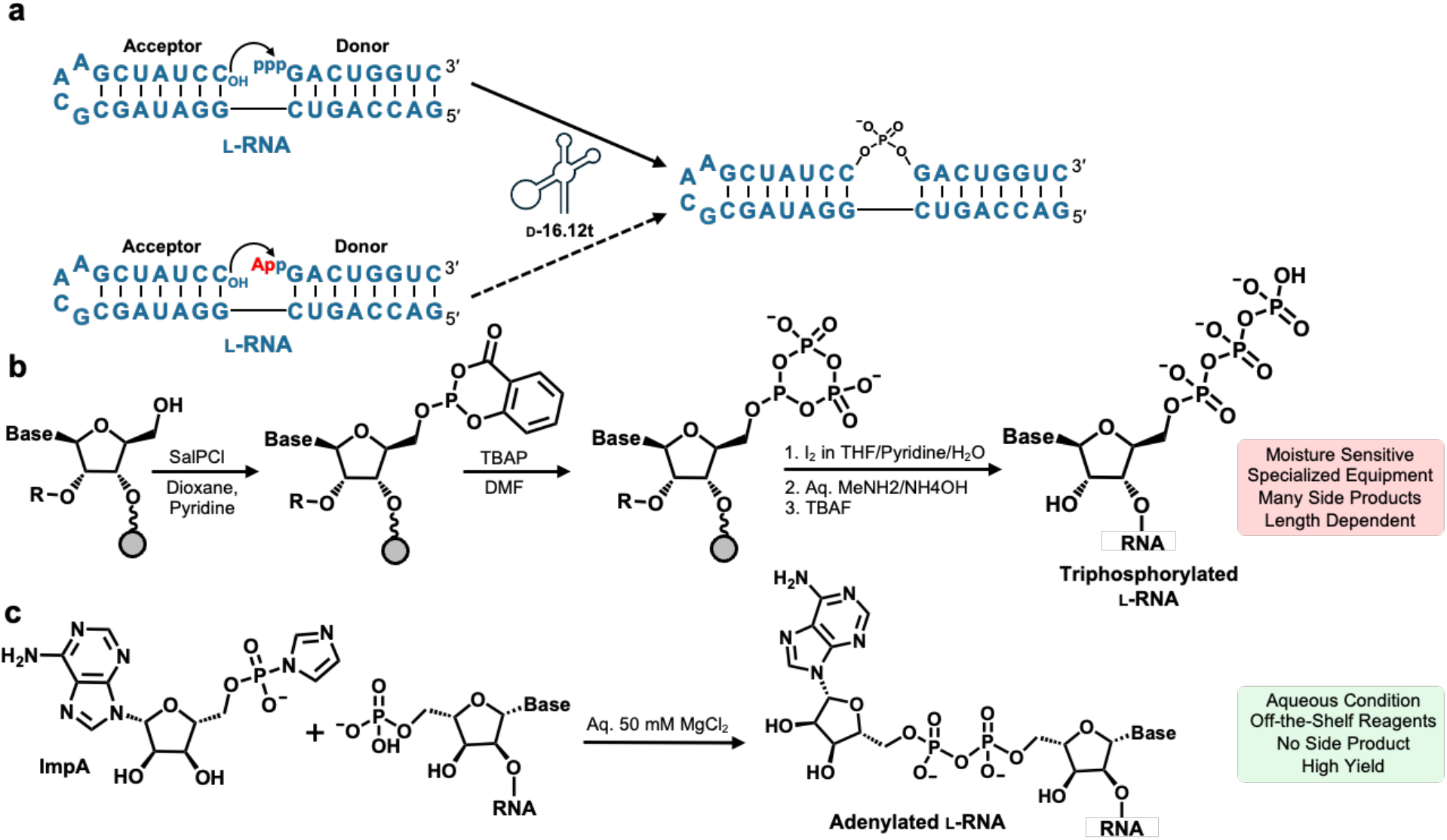
Cross-Chiral Ligation of L-RNA. (a) The D-16.12t ribozyme catalyzes ligation of the 3′-hydroxyl (OH) of the “acceptor” L-RNA substrate to the 5′-triphosphate (ppp) of the “donor” L-RNA substrate, resulting in formation of a 3′,5′-phosphodiester linkage (solid arrow). Whether this reaction occurs with 5′-adenylated (App) or related substrates is the subject of this study (dotted arrow). (b) Scheme for the chemical 5′-triphosphorylation of synthetic oligonucleotides using Ludwig-Eckstein chemistry. (c) Scheme for the chemical adenylation of synthetic oligonucleotides using adenosine-5′-phosphoimidazolide (ImpA). SalPCl = salicyl phosphorochloridite; TBAP = tributylammonium pyrophosphate; THF = tetrahydrofuran; TBAF = tetrabutylammonium fluoride.

Compared to chemical 5′-triphosphorylation, 5′-adenylation of ONs is synthetically more tractable.^39-41^ For example, 5′-adenylated ONs can be readily prepared from synthetic 5′-phosphorylated ONs by incubation with an aqueous mixture of adenosine-5′-phosphoimidazolide (ImpA) and MgCl_2_ (Figure 1c).^39^ This reaction is not moisture sensitive and, thus, does not require special equipment or carefully reagent handling to maintain anhydrous conditions. More importantly, this reaction is high-yielding and does not form difficult to remove side products. Recently, Höbartner and colleagues reported an RNA ligase DNAzyme that utilize 5′-adenylated RNAs as donor substrates and demonstrated that its ligase activity was equivalent to DNAzymes that utilize 5′-triphosphorylated donors.^42^ With this in mind, the goal of this study was to determine whether previously reported cross-chiral ligase ribozymes would accept 5′-adenylated L-RNA donors (or analogues thereof) prepared using the more easily accessible phosphoimidazolide chemistry and, if so, to assess the overall practicality of this approach for assembling long L-RNA molecules.

## RESULTS AND DISCUSSION

### Synthesis of 5′-adenylated L-RNAs

We first set out to prepare 5′-adenylated donor substrates for the ligation experiments, as well as demonstrate the compatibility of this charging reaction with long L-RNAs. Our initial substrate was an 8-mer L-RNA (L-RNA_8_; Figure 2a) that serves as the donor for the ligation reaction depicted in Figure 1a. This ligation complex is identical to the one originally used to evolve the 16.12t cross-chiral ligase.^25^ 5′-adenylated L-RNA_8_ (L-AppRNA_8_) was prepared from synthetic 5′-phosphorylated L-RNA_8_ by incubation with ImpA and MgCl_2_, as previously described (Figure 1c).^39^ As shown in Figure 2b,c, the adenylation reaction proceeded to nearly 80% completion after 5 hours and the 5′-adenylated product L-AppRNA_8_ was readily purified to >90% via polyacrylamide gel electrophoresis (PAGE) (Figure S1a). While this result was encouraging, efficient assembly of long L-RNAs by cross-chiral ligation will require the use of much longer substrates, which have proven challenging to activate via 5′-triphosphorylation using Ludwig-Eckstein chemistry.^33, 35^ Therefore, we proceeded to examine the efficiency of the adenylation reaction using progressively longer 5′-monophosphorylated L-RNAs (Figure 2a). We note that each substrate was a 3′-extension of L-RNA_8_, allowing them to be used in the same ligation complex (Figure 1a). To our surprise, all L-RNA donors could be charge to at least 50%, including the 50-mer L-pRNA_50_ (55%) (Figure 2c and Figure S1b-d). This indicated that the 5′-adenylation reaction provides an efficient approach for generating L-RNA donors for the eventual assembly of long L-RNAs by cross-chiral ligation.

**Figure 2.**
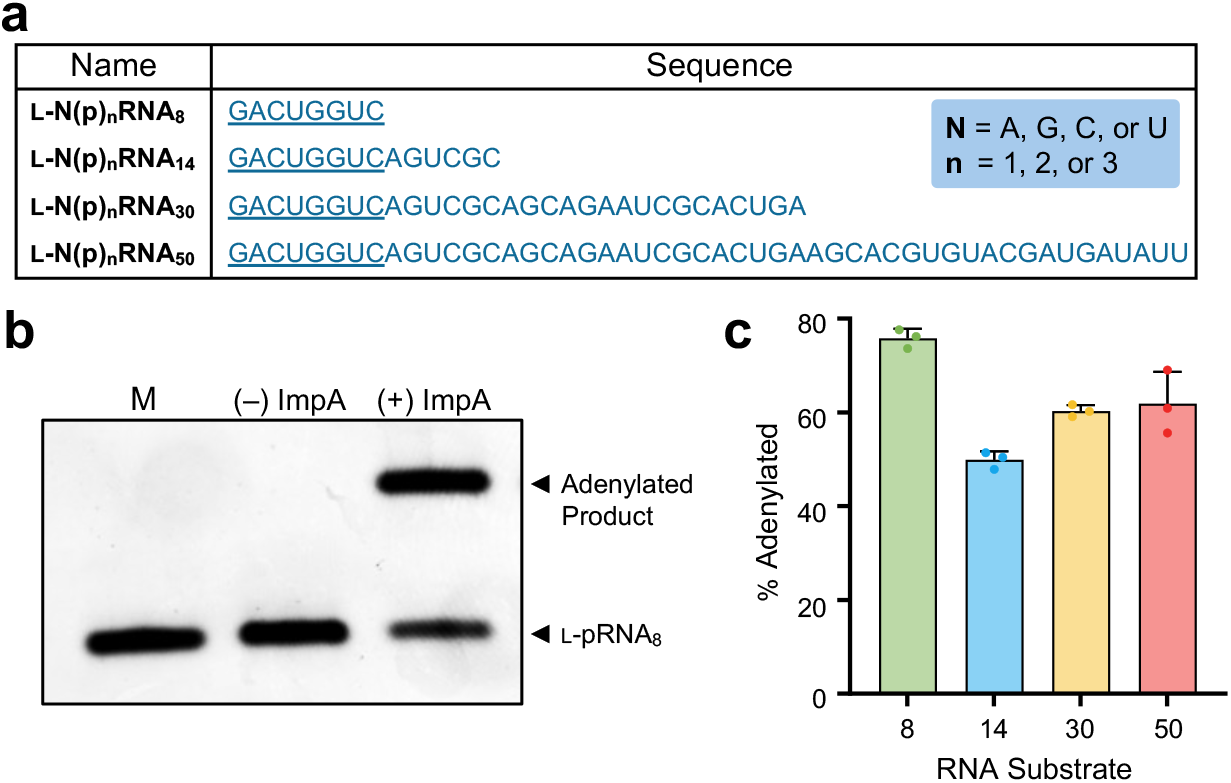
5′-Adenylation of L-RNA. (a) Sequences of the L-RNA donors used in this study (5′→3′). Underlined text indicates sequence complementarity with the hairpin acceptor depicted in Figure 1a. A complete list of oligonucleotides used in this study can be found in Table S1. (b) Representative denaturing PAGE analysis of the 5′-adenylation reaction. L-pRNA_8_ (100 µM) was incubated with or without ImpA (100 mM) in a reaction buffer containing 50 mM MgCl_2_ at 52 °C for 5 h. (c) Adenylation yields as determined by gel electrophoresis for different length L-RNAs donors. Error bars show standard deviation (n = 3).

### Cross-chiral ligase 16.12t is compatible with 5′-adenylated donors

With the 5′-adenylated L-RNA donors in hand, we investigated their compatibility with the cross-chiral ligation reaction catalyzed by D-16.12t. The 8-mer L-AppRNA_8_ donor was assembled into the corresponding substrate complex (Figure 1a) and treated with excess D-16.12t ribozyme. As controls, we also performed the reaction using 5′-phosphorylated and 5′-triphosphorylated versions of the same donor (L-pRNA_8_ and L-pppRNA_8_, respectively). As shown in Figure 3a, D-16-12t was indeed capable of catalyzing the ligation of the 5′-adenylated donor L-AppRNA_8_, yielding ∼30% ligated L-RNA product after 24 hours. Mass spectrometry (MS) analysis of the purified product confirmed that it contained the expected monophosphate linkage (Figure S2). For comparison, the reaction employing the 5′-triphosphorylated substrate L-pppRNA_8_ resulted in >50% ligated product after just 10 hours. The reduced activity observed for L-AppRNA_8_ was not unexpected, as the D-16.12t ribozyme was evolved to recognize and ligate 5′-triphosphorylated L-RNA. Furthermore, assuming a similar mechanism between the two substrates (i.e., S_N_2 at the α phosphate), the pyrophosphate leaving group associated with L-pppRNA_8_ is highly favored in the reaction compared to the adenosine monophosphate (AMP) leaving group associated with L-AppRNA_8_.^43^ No ligated product was observed for up to 24 hours when the 5′-phosphorylated donor (L-pRNA_8_) was used, and this reaction was indistinguishable from the no-ribozyme control (Figure 3a). The longer 5′-adenylated L-RNA donors also proved to be compatible with D-16-12t-mediated cross-chiral ligation, with product yields ranging between 20–40% after 24 hours (Figure 3b). As each of these donors harbored the same ligation junction and surrounding nucleotide sequence (Figure 2a), differences in ligation efficiency were likely due to the nature of the 3′-extension. Overall, these results demonstrated that 5′-adenylated L-RNA donors are compatible with the cross-chiral ligation reaction catalyzed by D-16.12t and can support the cross-chiral ligation of long L-RNA fragments.

**Figure 3.**
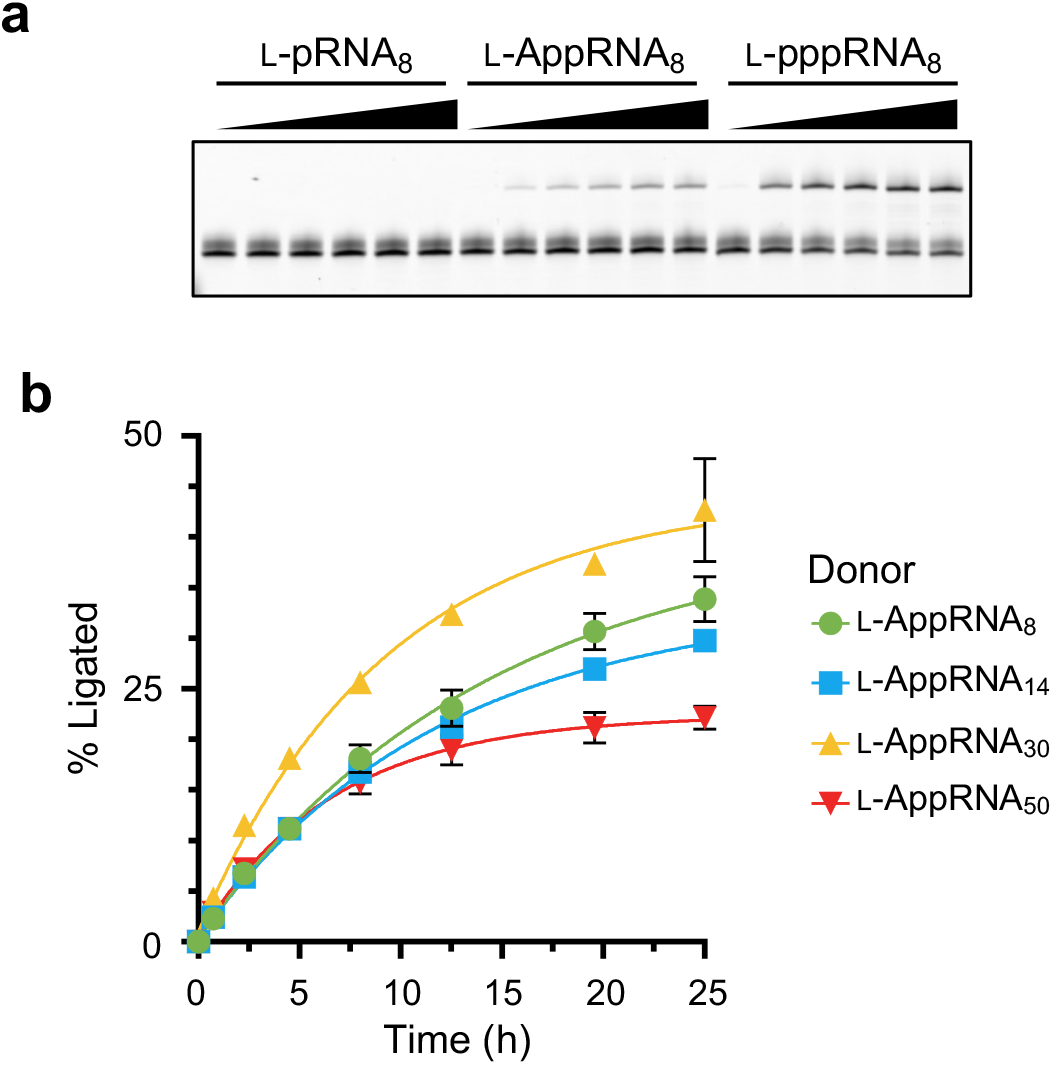
Cross-chiral ligation of adenylated L-RNA by D-16.12t. (a) Representative denaturing PAGE analysis for cross-chiral ligation of the indicated donor substrate by D-16.12t. A 5′-FAM-labeled version of the acceptor depicted in Figure 1a was used in these experiments. Reactions employed 10 μM D-16.12t, 1 μM acceptor substrate, 2 μM donor substrate, 250 mM NaCl, 250 mM MgCl_2_, and 50 mM Tris (pH 8.5) and were incubated at 23 °C for 2, 6, 10, 18, and 24 h. (b) Kinetic time course of D-16.12t ligating the indicated donor substrate. Reaction conditions are the same as in panel a. Error bars show standard deviation (n = 3).

### Cross-chiral ligation of 5′-adenylated donors is dependent on the identity of the β-phosphate nucleoside

Compared to an optimal 5′-triphosphorylated donor, 5′-adenylated donors contain a substitution of the γ-phosphate with an adenosine nucleoside. The possibility that 16.12t interacts directly with the γ-phosphate position prompted us to evaluate what role, if any, the identity of the substituted nucleoside plays in the reaction. Therefore, using the corresponding imidazolides, we prepared 5′-cytidylated (L-CppRNA_8_), 5′-guanylated (L-GppRNA_8_), and 5′-uridylated (L-UppRNA_8_) versions of the 8-mer L-RNA donor and subjected these substrates to the D-16.12t-catalyzed cross-chiral ligation reaction. Surprisingly, compared to L-AppRNA_8_, the new L-RNA donors performed far worse in the reaction (Figure 4a). Notably, the reaction employing the 5′-uridylated L-RNA donor (L-UppRNA_8_) yielded <3% ligated product after 24 hours, which was ∼10-fold less than L-AppRNA_8_ during the same period. The purity of all 5′-N-ylated donors was determined to be >90% by MS (Figure S3), indicating that these results were not due to differences in substrate quality.

**Figure 4.**
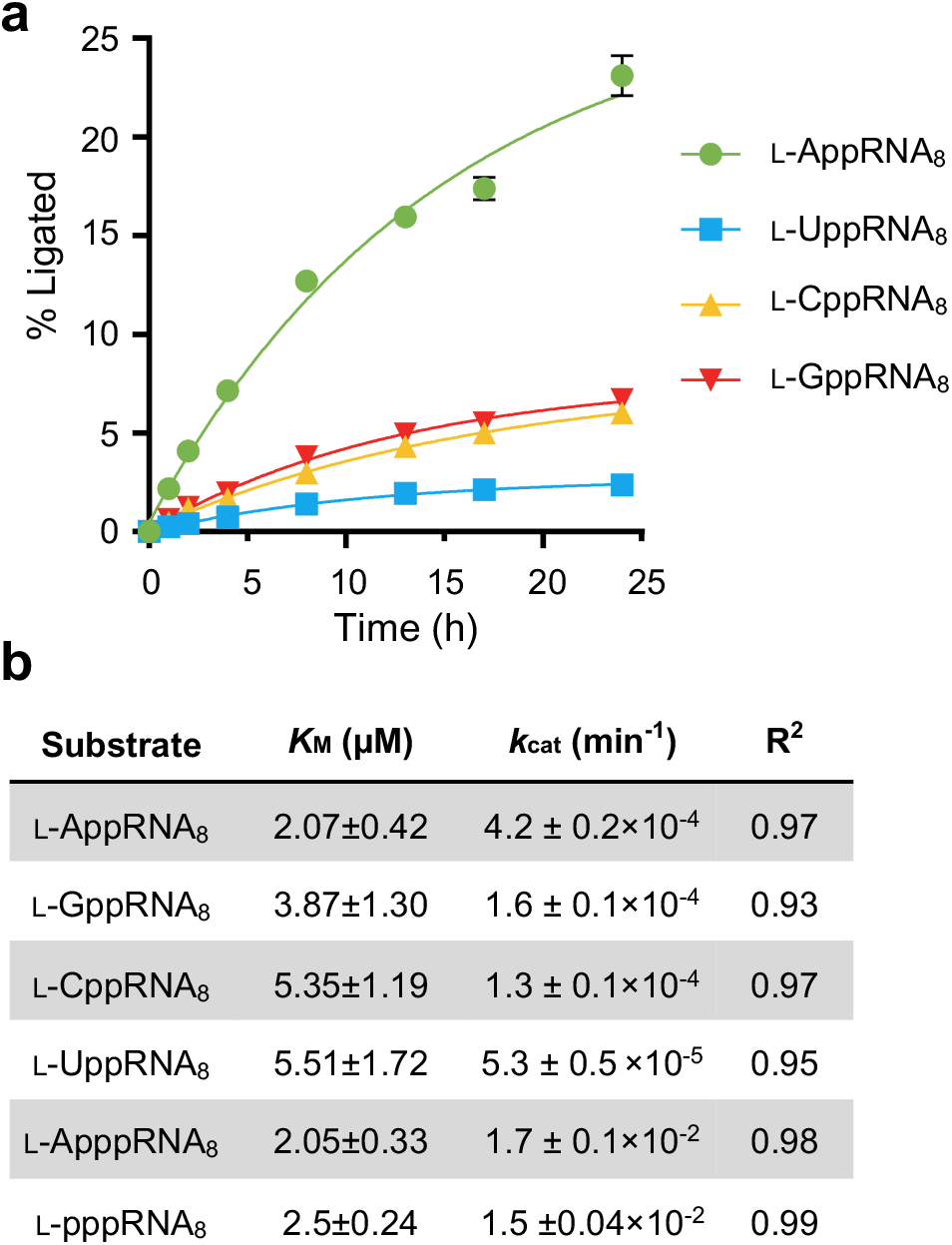
Cross-chiral ligation of 5′-adenylated donors is dependent on the identity of β-phosphate nucleoside. (a) Kinetic time course of D-16.12t ligating the indicated L-RNA donor substrate. Reaction conditions are the same as in Figure 3a. Error bars show standard deviation (n = 3). (b) Michaelis-Menten parameters for the ligation of the indicated L-RNA donor by D-16.12t. Data is mean ± standard deviation (n = 3).

To examine this phenomenon more carefully, we determined Michaelis-Menten constants for the ligation of the four different 5′-N-ylated L-RNA donors by D-16-12t (Figure S4). *K*_m_ and *k*_cat_ values are listed in Figure 4b and reveal a similar trend as the time-course experiments above. *K*_m_ values for the ligation of 5′-N-ylated donors were comparable to the ligation of the 5′-triphosphorylated donor (L-pppRNA_8_), increasing ∼2-fold for the poorest substrate (L-UppRNA_8_). However, *k*_cat_ values for the ligation of 5′-N-ylated donors decreased dramatically. For example, while the ligation reactions for L-AppRNA_8_ and L-pppRNA_8_ had nearly identical *K*_m_ values, the *k*_cat_ for ligation of L-AppRNA_8_ was ∼35-fold slower than L-pppRNA_8_. This strongly suggested that the reduced activity of D-16-12t with 5′-N-ylated L-RNA donors relative to the 5′-triphosphorylated donor is due to a much slower chemical step rather than weaker substrate binding. Again, this is consistent with the triphosphate being the more favored substrate in the S_N_2 reaction mechanism.^43^ Interestingly, the *k*_cat_ values among the four 5′-N-ylated donors differed by as much as 8-fold (Figure 4b), despite the reactions all taking place with the same diphosphate moiety. This indicated the involvement of the β-phosphate-linked nucleoside in the catalytic step, the identity of which either favors or disfavors the reaction, possibly due to unique interactions of the nucleobases with the ribozyme. Future studies will be needed to determine the exact mechanism(s) underlying these observations. Taken together, these data show that D-16-12t-mediated cross-chiral ligation of 5′-N-ylated L-RNA donors is dependent on the identity of the β-phosphate-linked nucleoside, with 5′-adenylated substrates being highly preferred in the reaction.

### Cross-chiral ligation is compatible with 5′-adenosyl triphosphate L-RNA

Given the efficiency by which adenylated substrates could be produced from ImpA, we investigated whether a similar approach could be used to generate L-RNA donors with a 5′-triphosphate moiety, which were expected to be a better substrates for cross-chiral ligation. To begin, we prepared adenosine-5′-diphosphate (ADP)-imidazolide (ImppA) (Figure 5a) and subjected it to the same coupling reaction with L-pRNA_8_ and L-pRNA_14_ to generate the corresponding 5′-adenosyl triphosphorylated L-RNAs. Crude reaction mixtures were analyzed by PAGE (Figure S5a) and MS (Figure 5b and Figure S5), revealing that L-pRNA_8_ and L-pRNA_14_ had undergone >80% conversion to the corresponding 5′-adenosyl triphosphates (L-ApppRNA_8_ and L-ApppRNA_14_, respectively). Importantly, little to no undesirable side-products were generated during the reaction. For comparison, we also attempted to prepare L-pppRNA_14_ by chemical 5′-triphosphorylation using Ludwig-Eckstein chemistry (Figure 1b) following the rigorous procedures established by Bare et al.^38^ In this case, a large amount of side products were observed (Figure 5b), including the 5′-monophosphate, 5′-H-phosphonate, and the 5′-diphosphate, which are commonly produced during this reaction and are very challenging to separate from the desired product, especially for longer oligonucleotides.^33, 38^ Indeed, attempts to purify L-pppRNA_14_ by PAGE were unsuccessful at completely removing these impurities (Figure S6a,b). Furthermore, HPLC-purified L-pppRNA_8_ obtained from a commercial source was only ∼70% pure and still contained various hypo-phosphorylated impurities (Figure S6c). In contrast, after PAGE purification, L-ApppRNA_14_ generated from ADP-imidazolide was >90% pure and contained none of the aforementioned side products (Figure S5). Taken together, these results show that ADP-imidazolide provides a practical, fast, and high-yielding approach to generate high purity 5′-adenosyl triphosphorylated oligonucleotides and is attractive alternative to the more arduous Ludwig-Eckstein chemistry.

**Figure 5.**
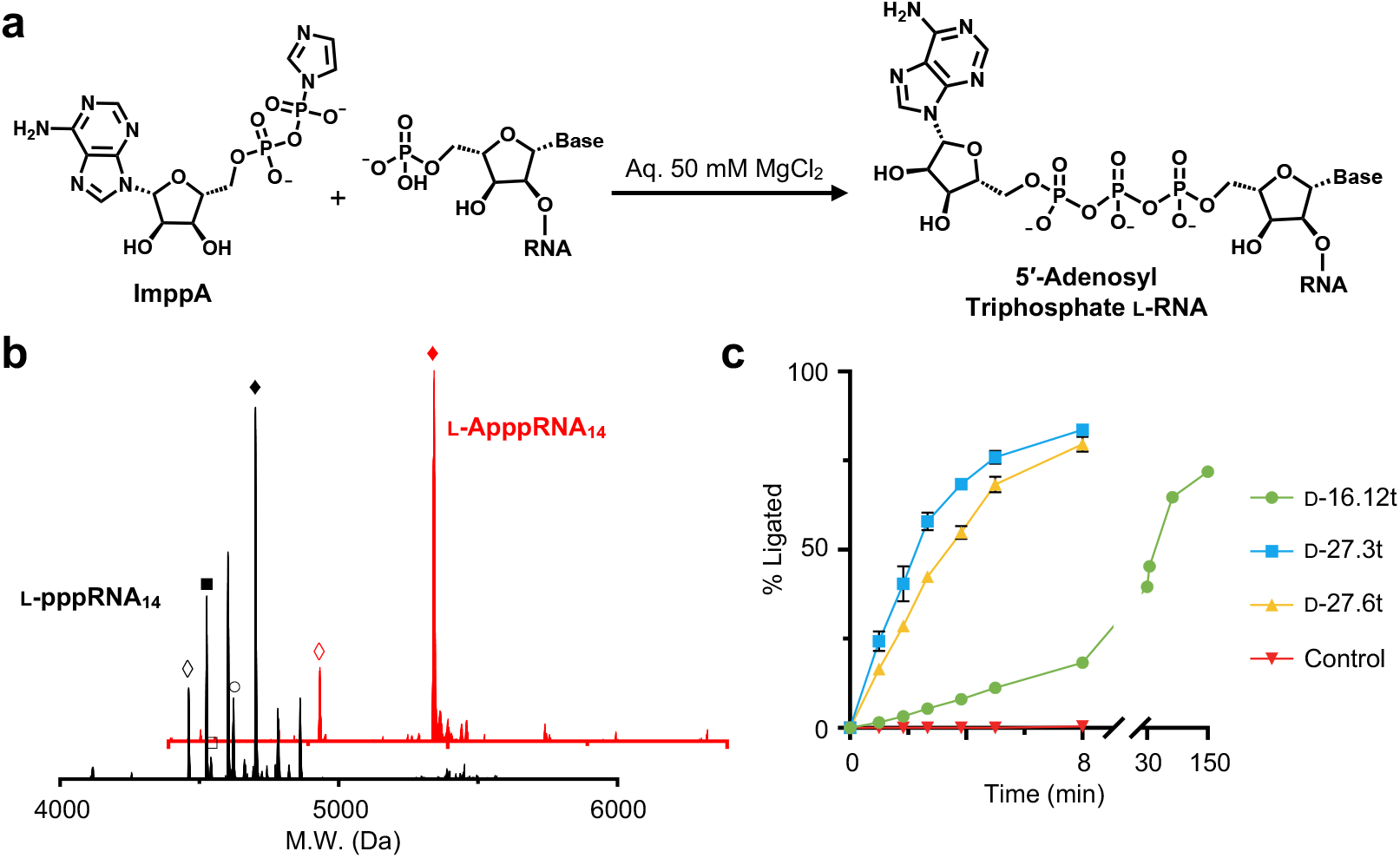
Cross-chiral ligation is compatible with 5′-adenosyl triphosphates. (a) Scheme for the chemical activation of synthetic oligonucleotides using adenosine-5′-diphosphate (ADP)-imidazolide (ImppA). (b) Representative mass spectrometry analysis of crude products from (i) the reaction of L-pRNA_14_ with ImppA (red) or (ii) the chemical triphosphorylation of L-RNA_14_ via Ludwig-Eckstein chemistry (black). Open diamond = starting material; closed diamond = desired product; open square = 5′-monophosphate; closed square = 5′-H-phosphonate; open circle = 5′-diphosphate. (c) Kinetic time course of the indicated D-RNA ribozyme ligating L-ApppRNA_8_ to the acceptor depicted in Figure 1a. The control experiment lacks a ribozyme. Reaction conditions are the same as in Figure 3a. Error bars show standard deviation (n = 3).

We next tested the compatibility of L-ApppRNA_8_ with cross-chiral ligation by D-16.12t. As shown in Figure 5c, the ligation reaction was ∼75% complete after 2.5 hours, marking a dramatic improvement compared to reactions employing the 5′-N-ylated donors (e.g., L-AppRNA_8_) (Figure 4a). Mechaelis-Menten constants for the ligation of L-ApppRNA_8_ were almost identical to L-pppRNA_8_, indicating that the adenosine moiety had little impact on the ligation reaction when linked to the γ-phosphate (Figure 4b). Furthermore, MS analysis confirmed that the ligated products generated using either L-ApppRNA_8_ or L-pppRNA_8_ as donors were identical (Figure S7). In addition to 16.12t, we also examined two other cross-chiral ligases ribozymes, namely 27.3t and 27.6t (Table S1), for their compatibility with this new class of donor substrate. These ribozymes were evolved from 16.12t for templated ligation of 5′-triphosphorylate trinucleotides and show greatly improved ligation activity (10^2^–10^3^-fold) compared to the parental ribozyme.^26^ Indeed, cross-chiral ligation reactions employing these ribozymes and the donor L-ApppRNA_8_ were nearly complete in under 10 minutes (Figure 5c). Thus, the combination of 5′-adenosyl triphosphate donors and cross-chiral ligases 27.3t or 27.6t provides a promising strategy for ligating L-RNAs and, potentially, for assembling large L-RNAs from multiple components.

### 5′-Adenosyl triphosphates are compatible with large L-RNA assembly

The key motivation behind this work was the development of a more efficient strategy for assembling long L-RNAs by cross-chiral ligation. Therefore, having shown the ease by which compatible 5′-triphosphorylated L-RNA donors could be produced using ImppA and the efficient ligation of these substrates by D-27.3t and D-27.6t, we attempted to assemble a large L-RNA using this overall approach. For this study, we chose to assemble the L-RNA version of the 27.6t ribozyme (Figure 6a), which at 129 nt long, is well beyond the limit of current solid-phase synthesis methods. Moreover, we sought to use the newly assembled L-27.6t ribozyme to ligate 5′-adenylated D-RNAs, demonstrating the compatibility of our approach with both stereochemical orientations of the system. The ribozyme was broken up into two L-RNA fragments: a 64-nt acceptor (L-27.6t_A_) and a 65-nt donor (L-p27.6t_D_) (Figure 6a). Following solid-phase synthesis, the 5′-monophosphorylated donor was reacted with ImppA as before to generate the corresponding 5′-adenosyl triphosphate (L-Appp27.6t_D_). Analysis of the crude reaction mixture revealed nearly complete conversion of L-p27.6t_D_ into L-Appp27.6t_D_ (>95%; Figure S12), which was used without further purification. To the best of our knowledge, this is the longest L-RNA produced synthetically containing a 5′-triphosphate moiety. When combined with the splint (L-27.6_s_; Table S1), cross-chiral ligation of L-27.6t_A_ and L-Appp27.6t_D_ by D-27.3t proceeded to ∼60% yield (35% isolated yield) after only 3 hours (Figure 6b). MS analysis of the PAGE purified product confirmed the correct assembly of L-27.6t (Figure S8). Importantly, the newly assembled L-27.6t was functional (Figure 6c). Addition of L-27.6t to the D-RNA substrate complex shown in Figure S9a resulted in the formation of >80% ligated D-RNA product in less than 10 minutes, which was similar to the D-ribozyme (prepared by *in vitro* transcription) acting on an L-RNA substrate complex (Figure 6c). Thus, both reaction configurations (D-ribozyme and L-substates versus L-ribozyme and D-substrates) are compatible with 5′-adenosyl triphosphate donors. Moreover, as both D- and L-RNA donors were generated using ImppA containing a D-ribose moiety, the stereochemistry of the γ-phosphate-linked nucleoside relative to the RNA substrates and ribozyme does not appear to impact the reaction. Finally, the product of the ligation of the two D-RNA substrates was subjected to RNase A digestion, which only cleaves 3′,5′-phosphodiester linkages. Complete cleavage was observed at the ligation junction (Figure S9b), indicating that cross-chiral ligation using donors activated with 5′-adenosyl triphosphates produces natural 3′,5′-phosphodiester linkages.

**Figure 6.**
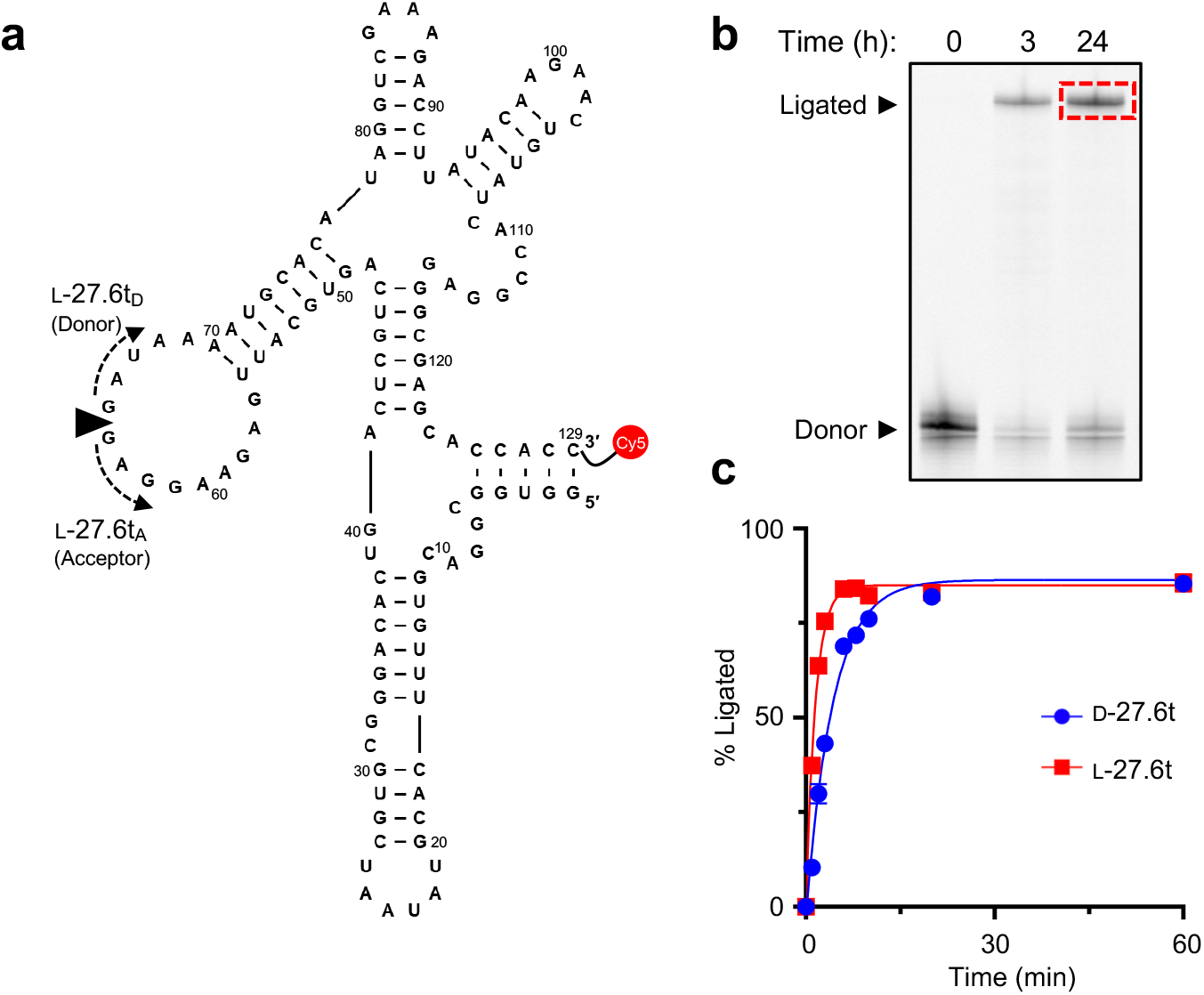
Assembly and functional analysis of L-27.6t. (a) Sequence and secondary structure of the 27.6t ribozyme. Black wedge indicates the ligation junction, with the donor and acceptor sequences on either side. (b) Representative denaturing PAGE analysis for cross-chiral assembly of L-27.6t. Reactions employed 5 μM D-27.3t, 2.5 μM acceptor substrate, 2.5 μM donor substrate, 2.5 μM splint strand, 250 mM NaCl, 250 mM MgCl_2,_ and 50 mM Tris (pH=8.5), which were incubated for the indicated times at 23 °C. The red box indicates the ligated product that was excised from the gel for further analysis. (c) Catalytic activity of L-27.6t assembled by cross-chiral ligation. For comparison, the ligation of L-RNA by D-27.6t prepared by *in vitro* transcription is also shown. The reaction conditions are the same as in Figure 3a, except that 5 μM ribozyme was used. The substrates for this reaction are depicted in Figure S9a. Error bars show standard deviation (n = 3).

## CONCLUSIONS

In summary, this work expands the substrate scope of cross-chiral ligase ribozymes to include 5′-adenylated (or related) L-RNAs as donors, opening the door to a straightforward and practical approach for assembling long L-RNA. Until now, a key bottleneck in the cross-chiral assembly of long L-RNA molecules from multiple shorter fragments has been obtaining 5′-triphosphorylated L-RNA donors in sufficient purity and quantity to support the ribozyme-catalyzed ligation reaction. Herein, we show that compatible donors can be readily obtained by coupling 5′-phosphorylated L-RNA with either ImpA or ImppA, the latter yielding 5′-adenosyl triphosphate donors having activity equivalent to the native 5′-triphosphate. Compared to other chemical triphosphorylation approaches (e.g., Ludwig-Eckstein chemistry^36^), the advantages of this method are: 1) The procedure is straightforward and does not require careful reagent preparation or specialized equipment; 2) The yields are high, even for long (>50-nt) L-RNAs; and 3) It does not generate difficult to purify side products. To demonstrate the utility of this approach for assembling long L-RNA molecules, we synthesized a 65-nt L-RNA containing a 5′-adenosyl triphosphate in high yield and purity, which was subsequently assembled into the L-RNA version of the 129-nt 27.6t ribozyme via cross-chiral ligation. To the best of our knowledge, this is the longest L-RNA molecules assemble by a single ligation event to date. We anticipate that further optimization will lead to even more efficient donor activation chemistries and L-RNA assembly strategies. For example, we showed that the identity of the β-phosphate-linked nucleoside greatly impacted cross-chiral ligation of the 5′-adenylated donors (Figure 4a), possibly as a result of direct interactions with the ribozyme. Thus, screening a larger repertoire of chemically modified nucleosides harboring different classes of functional groups (e.g., cations, anions, polar groups, etc.) may yield donors with improved ligation activity. Further *in vitro* evolution of the cross-chiral ribozymes employing 5′-adenosyl L-RNA donors (both di- and triphosphates) is also expected to produce more efficient catalysts and warrants future investigation.^25, 26^

Overall, given the practical advantages of phosphoimidazolide chemistry for preparing compatible 5′-triphosphorylated L-RNAs, coupled with the ready availability of D-cross-chiral ribozymes via standard *in vitro* transcription reactions, we expect that the methods reported herein will make long L-RNAs more accessible to a wider range of researchers and will enable a variety of practical applications. For example, because this approach has few sequence or other design constraints, it could be used to generate nuclease-resistant, mirror-image versions of many RNA-based molecular sensors and other devices, thereby greatly expanding the utility of these technologies for *in vivo* applications. Moreover, the ability to easily generate long L-RNAs will further support ongoing efforts to build a mirror-image biology systems, such as mirror-image ribosome that translates D-proteins. Finally, although this work focused on the chemical activation and ligation of L-RNA, our straightforward method for generating γ-substituted 5′-triphosphates can be readily applied to any nucleic acid polymer, providing researchers access to ONs harboring a diverse range of triphosphate analogues and mRNA 5′-cap modifications.

## ASSOCIATED CONTENT

### Supporting Information

Reagents and experimental methods; supplementary figures and tables; mass spectrometry characterization of oligonucleotides; oligonucleotides sequences and structures; results from RNase A digestion.

## AUTHOR INFORMATION

### Author contributions

Xuan Han: Methodology, Investigation, Formal analysis, Visualization, Writing – Review & Editing. Jonathan T. Sczepanski: Conceptualization, Writing – Original Draft, Supervision, Project administration, Funding acquisition.

### Notes

The authors declare no competing financial interests.

## ACKNOWLEDGMENT

This work was supported by the National Science Foundation (2114588) and the National Institute of General Medical Sciences (R35GM124974) of the National Institutes of Health. The content is solely the responsibility of the authors and does not necessarily represent the official views of the National Institutes of Health.

