## Supplemental Information for "An Expanded Substrate Scope for Cross-Chiral Ligation Enables Efficient Synthesis of Long L-RNAs"

### S1. Supplementary Text.

### MATERIALS AND METHODS

#### General

All DNA oligonucleotides were purchased from Integrated DNA Technologies (Coralville, IA). All RNA oligonucleotides were prepared by solid-phase oligonucleotide synthesis using an Expedite 8909 DNA Synthesizers, except for L-pppRNA$_8$-CG, which was purchased from ChemGenes (Wilmington, MA). D-RNA phosphoramidites, solid supports, and other oligonucleotide synthesis reagents were purchased from Glen Research (Sterling, VA). L-Nucleoside phosphoramidites and solid supports were purchased from ChemGenes (Wilmington, MA). Superscript II reverse transcriptase and RNAse I was purchased from ThermoFisher (Waltham, MA). Recombinant T7-RNA polymerase was expressed in house as previously described.[1] RNase A was purchased from New England Biolabs (Ipswich, MA). Sulfo-Cy3 N-Hydroxysuccinimide (NHS) ester and Sulfo-Cy5 NHS esters were purchased from Lumiprobe (Cockeysville, MD). All other reagents were purchased from Sigma Aldrich (St. Louis, MO).

#### Oligonucleotide synthesis and purification

All oligonucleotides used in this study are shown in Table S1. All L-RNAs were synthesized in house on 8909 DNA synthesizer using manufacturer recommended protocols. 5′-Fluorescein labeling of acceptor strands was accomplished using the 6-FAM phosphoramidite (Glen Research, Sterling, VA). The 3′-biotin modification was installed using the 3′-biotinTEG CPG (Glen Research, Sterling, VA). Sulfo-Cy3/5 NHS esters were conjugated to the 3′ end of the indicated L-RNAs via a 3′ amino modification installed at the time of synthesis. The conjugation reaction has been described in our previous work.[2] Prior to use, all synthetic oligonucleotides were purified by 20% denaturing polyacrylamide gel electrophoresis (PAGE; 19:1, acrylamide/bisacrylamide). Pure oligonucleotides were excised from the gel and eluted overnight at room temperature in a buffer consisting of 10 mM Tris (pH 7.6), 200 mM NaCl, and 10 mM EDTA. The filtered supernatants were concentrated using either a 3 kDa pore size Amicon Ultra Centrifugal Filter device (MilliporeSigma, Burlington, MA) or a SepPak C18 cartridge (Waters, Milford, MA) following the manufacturer's recommended procedures. All purified oligonucleotides were desalted by ethanol precipitation prior to use. Oligonucleotide concentrations were determined by absorbance at 260 nm on a NanoDrop 2000c (ThermoFisher, Waltham, MA) and the identity of all oligonucleotides was confirmed using a Thermo Scientific Q Exactive Focus ESI mass spectrometer. Deconvoluted mass spectrometry data was generated using UniDec 7.0.2.[3] Oligonucleotide purity was determined by ESI-MS using relative peak intensities.

#### *In vitro* transcription of RNA

DNA templates used to prepare cross-chiral ribozymes D-16.12t, D-27.3t, and D-27.6t were generated by a cross-extension of two overlapping synthetic oligonucleotides (Table S2). The pair of overlapping oligonucleotides (200 pmol each) was added to 60 µL water and the solution was heated at 70 °C for 2 minutes followed by slow cooling to room temperature. The cooled solution was supplemented with reagents to achieve a final volume of 100 µL containing 3 mM $MgCl_2$, 75 mM KCl, 10 mM dithiothreitol (DTT), 50 mM Tris (pH 8.3), and 0.5 mM each of the four dNTPs. 10 U/µL Superscript II reverse transcriptase was added to initiate the extension reaction, which was allowed to proceed at 42 °C for 45 minutes. The DNA was then desalted via ethanol precipitation and the resulting pellet was directly resuspended in a transcription solution (1 mL) containing 25 mM $MgCl_2$, 2 mM spermidine, 10 mM DTT, 40 mM Tris (pH 7.9), and 5 mM of each of the four NTPs. The transcription reaction was initiated by the addition of 10 U/µL T7 RNA polymerase and 0.001 U/µL Inorganic Pyrophosphatase (IPP) and the mixture was incubated at 37 °C for 2 hours. At that point, 10 U TURBO DNase was added and the reaction mixture was incubated for an additional 30 minutes at 37 °C. The reaction mixture was then ethanol precipitated and the RNA was purified by denaturing PAGE (10%, 19:1 acrylamide/bisacrylamide) as described above.

#### Synthesis of phosphoimidazolides

Adenosine 5′-phosphoimidazolide (ImpA), guanosine 5′-phosphoimidazolide (ImpG), uridine 5′-phosphoimidazolide (ImpU), cytidine 5′-phosphoimidazolide (ImpC), and adenosine 5′-diphosphate (ADP)-imidazolide (ImppA) were prepared in the same manner using a procedure adapted from Hafner et al.[4] A solution containing 2,2’-dipyridyldisulfide (220 mg, 1 mmol) in 3 mL of dry DMF was added dropwise into a stirring solution (12 mL) of triphenylphosphine (262 mg, 1 mmol), imidazole (170 mg, 2.5 mmol), and triethylamine (0.9 mL, 2.5 mmol) in dry DMF under argon. A suspension of 0.5 mmol nucleotide 5′-mono- or 5′-diphosphate in 15 mL dry DMF was then added dropwise to the same stirring solution above. The reaction was stirred for an additional 5 hours under argon. The completed reaction was precipitated via the dropwise addition to a stirring solution of sodium perchlorate (1.1g, 9 mmol) in 110 mL of acetone and 55 mL of anhydrous diethyl ether. The precipitant was collected via centrifugation at 5000 rpm for 5 minutes then washed with acetone followed by diethyl ether. The crude phosphoimidazolide products were dried under vacuum and characterized by mass spectrometry. These reactions typically resulted in the conversion of >80% of the nucleotide into the corresponding phosphoimidazolide (Figure S15), and the crude product was used without further purification.

**General procedure of the synthesis of 5′-N-ylated RNAs**

5′-monophosphorylated RNAs (5 nmol) were dissolved in 50 µL of an aqueous solution containing 50 mM $MgCl_2$ and 100 mM crude phosphoimidazolide product (based on theoretical yield). The reaction was incubated at 52 °C for 5 hours with an additional 25 µL of 100 mM phosphoimidazolide added at the 3 hour mark. To monitor the progress of the reaction, an aliquot was taken and resolved by denaturing PAGE (20%, 19:1 acrylamide:bis-acrylamide) and visualized by staining with 1 x SYBR Gold DNA stain (ThermoFisher, Waltham, MA). Completed reactions were then desalted using Amicon Ultra Centrifugal Filter MWCO 3k (Millipore-Sigma, Burlington, MA) and the oligonucleotide products were purified by denaturing PAGE as described above.

**Analysis of cross-chiral ligation activity**

The components for cross-chiral ligation were assembled in a reaction mixture containing 2 µM of the L-RNA acceptor (L-$\text{Acceptor}_{hp}$ for the ligation complex depicted in Figure 1a), 4 µM of L-RNA donor, 20 µM of the D-RNA ribozyme (16.12t, 27.3t, or 27.6t), 250 mM NaCl, and 50 mM Tris (pH 8.5). The mixture was heated at 70 °C for 2 minutes and cooled down to room temperature slowly. The reaction was initiated by the addition of an equal volume of a solution containing 500 mM $MgCl_2$, 250 mM NaCl, and 50 mM Tris (pH 8.5), and the reaction was incubated at 23 °C. Aliquots (2 µL) were taken at the indicated times and quenched with a solution (18 µL) containing 90% v/v formamide and 10 mM EDTA. The reaction products were then analyzed by denaturing PAGE (20%, 19:1 acrylamide/bisacrylamide). The gel was imaged by fluorescence emission (Cy5 excitation/emission: 635 nm/≥665 nm; Cy3 excitation/emission: 532 nm/≥575 nm; FAM excitation/emission: 473 nm/≥510 nm) using Typhoon FLA-9500 Multimode Molecular Imager and quantified using ImageQuant TL software (version 8.2) (General Electric Co., Boston, MA).

Michaelis–Menten parameters were determined using a similar reaction conditions as above, except that the concentration of the L-RNA ligation complex (L-$\text{Acceptor}_{hp}$ and indicated donor) was lowered to 0.05 µM and the concentrations of the D-RNA ribozyme was varied between 0 and 50 µM. Values for $k_{obs}$ were obtained for each enzyme concentration based on the initial rates, then fit to the Michaelis-Menten equation: $k_{obs} = k_{cat}[E] / (K_m + [E])$.[5]

**Cross-Chiral synthesis of L-27.6t ribozyme**

A 500 μL reaction mixture containing 5 μM L-27.6t$_A$, 5 μM L-Appp-27.6t$_D$, 5 μM L-27.6$_S$, 10 μM of D-RNA ligase ribozyme, 250 mM NaCl, and 50 mM Tris (pH 8.5) was heated to 70 °C for 2 minutes and cooled slowly to room temperature. The reaction was initiated by the addition of an equal volume (500 μL) of a solution containing 500 mM $MgCl_2$, 250 mM NaCl, and 50 mM Tris (pH 8.5) and was incubated at 23 °C for 5 hours. The reaction mixture was concentrated using a 3 kDa pore size Amicon Ultra Centrifugal Filter device and desalted by ethanol precipitation. The obtained pellet was resuspended in 50 μL of phosphate buffered saline PBS (137 mM NaCl, 10 mM phosphate, 2.7 mM KCl; pH 7.4) to which 50 U of RNase I was added to degrade the remaining D-RNA ribozyme. The mixture was incubated at 37 °C for 30 minutes, concentrated by ethanol precipitation and purified by denaturing PAGE (20%, 19:1 acrylamide/bisacrylamide) as described above.

**Determination of cross-chiral ligation regiospecificity**

The L-27.6t ribozyme was used to ligate a D-RNA substrate complex consisting of 20 μM D-RNA$_A$, 10 μM D-Appp-RNA$_D$, 20 μM D-RNA$_S$ (Table S1). The ligation reaction was performed as described above. The ligated product was purified by PAGE, desalted by ethanol precipitation, and dissolved in water. RNase A digestion was carried out in a reaction mixture containing 100 nM of the purified ligation product, 50 mM Tris (pH 7.6), and the indicated amounts of RNase A, which was incubated for 5 minutes at 23 °C. The reaction was quenched via the addition of 10 μg/μL tRNA and was analyzed immediately by denaturing PAGE (20%, 19:1 acrylamide/bisacrylamide) as described above. An authentic all-3′,5′-linked RNA (D-RNA$_{16}$) having an identical sequence to the ligated RNA product was treated with RNase A in a similar manner and used for comparison (Figure S9).[5]

**S2. Supplementary Figures.**

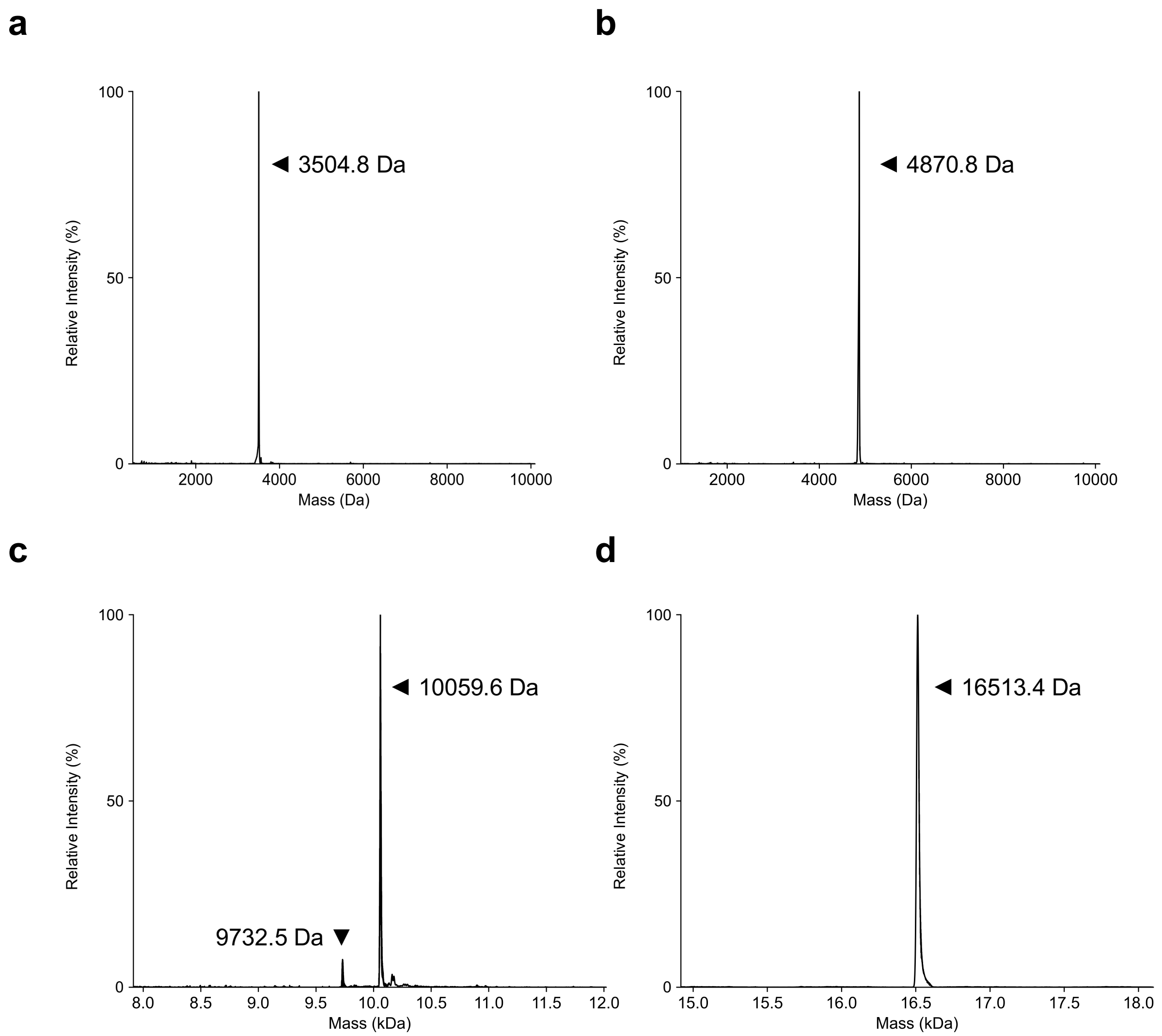


**Figure S1.** ESI-MS characterization of 5′-adenylated L-RNA donors. (a) ESI-MS data for L-AppRNA$_8$. Calculated: 3504.3 Da, observed: 3504.8 Da. (b) ESI-MS data for L-AppRNA$_{14}$. Calculated: 4870.8 Da, observed: 4870.8 Da. (c) ESI-MS data for L-AppRNA$_{30}$. Calculated: 10060.1 Da, observed: 10059.6 Da. (d) ESI-MS data for L-AppRNA$_{50}$. Calculated: 16513.8 Da, observed: 16513.4 Da.

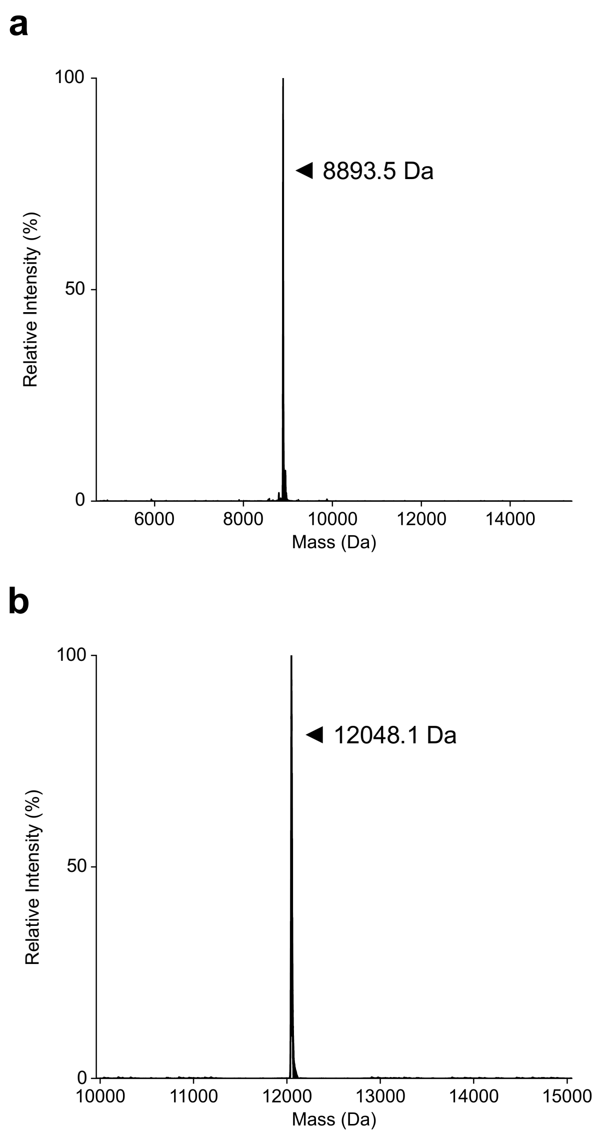


**Figure S2.** Cross-chiral ligation of L-Acceptor$_{hp}$ and L-AppRNA$_{8}$. (a) ESI-MS data for L-Acceptor$_{hp}$. Calculated: 8891.6 Da, Observed: 8893.5 Da. (b) ESI-MS data for ligated product of L-Acceptor$_{hp}$ and L-AppRNA$_{8}$. Calculated: 12049.4 Da, Observed: 12048.1.

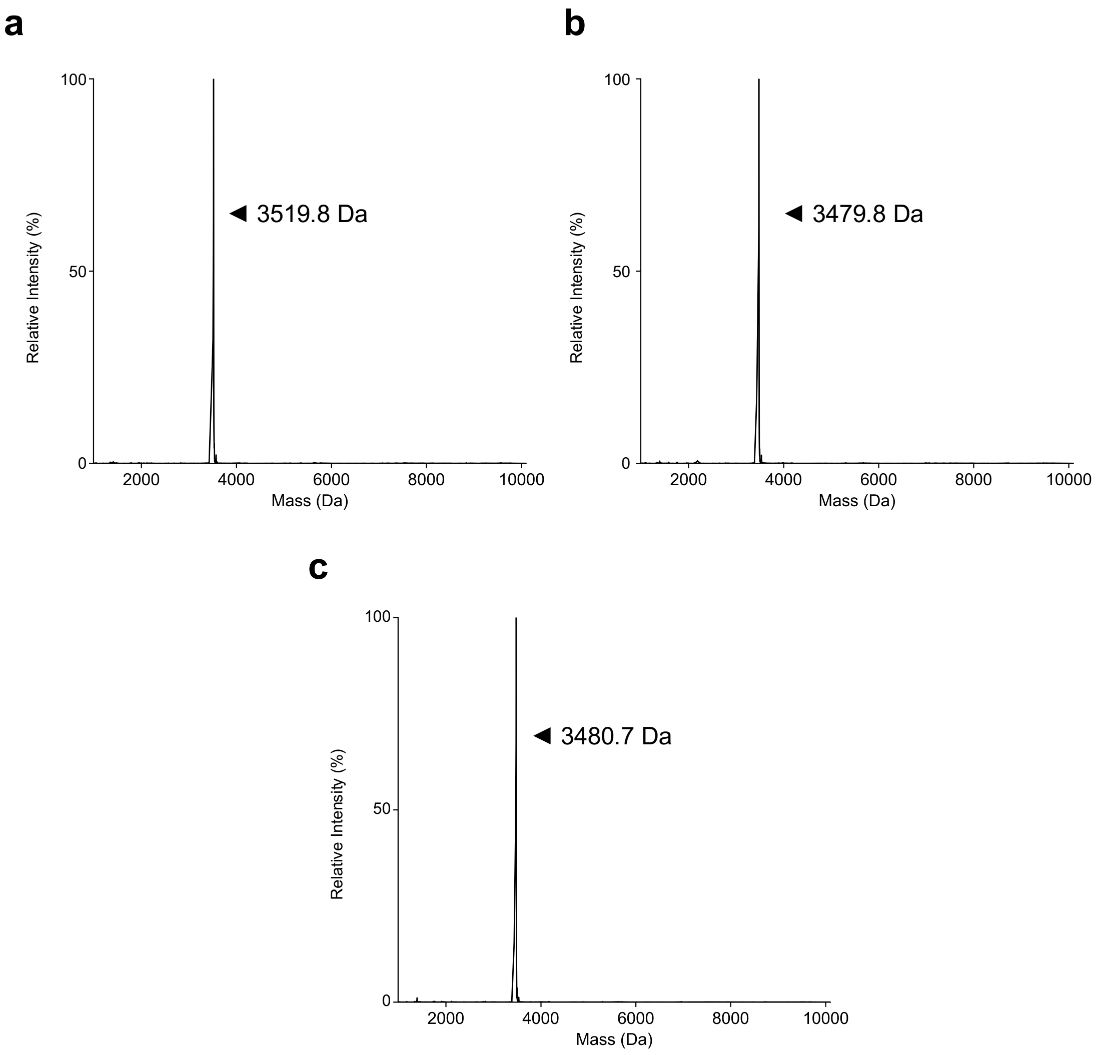


**Figure S3.** ESI-MS characterization of 5′-N-ylated L-RNA donors. (a) ESI-MS data for L-GppRNA$_8$. Calculated: 3520.3 Da, observed: 3519.8 Da. (b) ESI-MS data for L-CppRNA$_8$. Calculated: 3480.3 Da, observed: 3479.8 Da. (c) ESI-MS data for L-UppRNA$_8$. Calculated: 3481.3 Da, observed: 3480.7 Da.

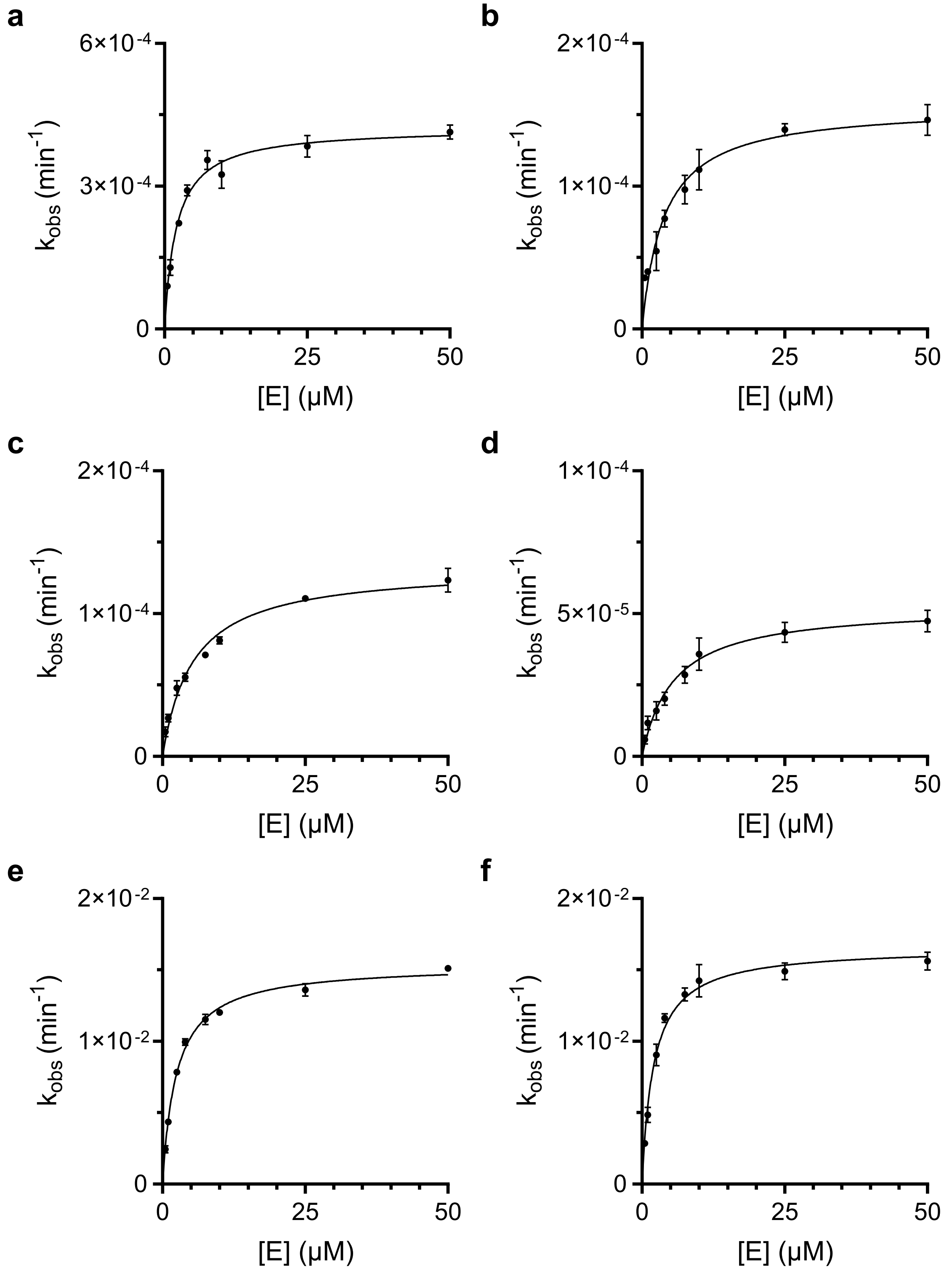


**Figure S4.** Fitted Michaelis-Menten kinetics analysis for (a) L-AppRNA$_8$, (b) L-GppRNA$_8$, (c) L-CppRNA$_8$, (d) L-UppRNA$_8$, (e) L-pppRNA$_8$, and (f) L-ApppRNA$_8$.

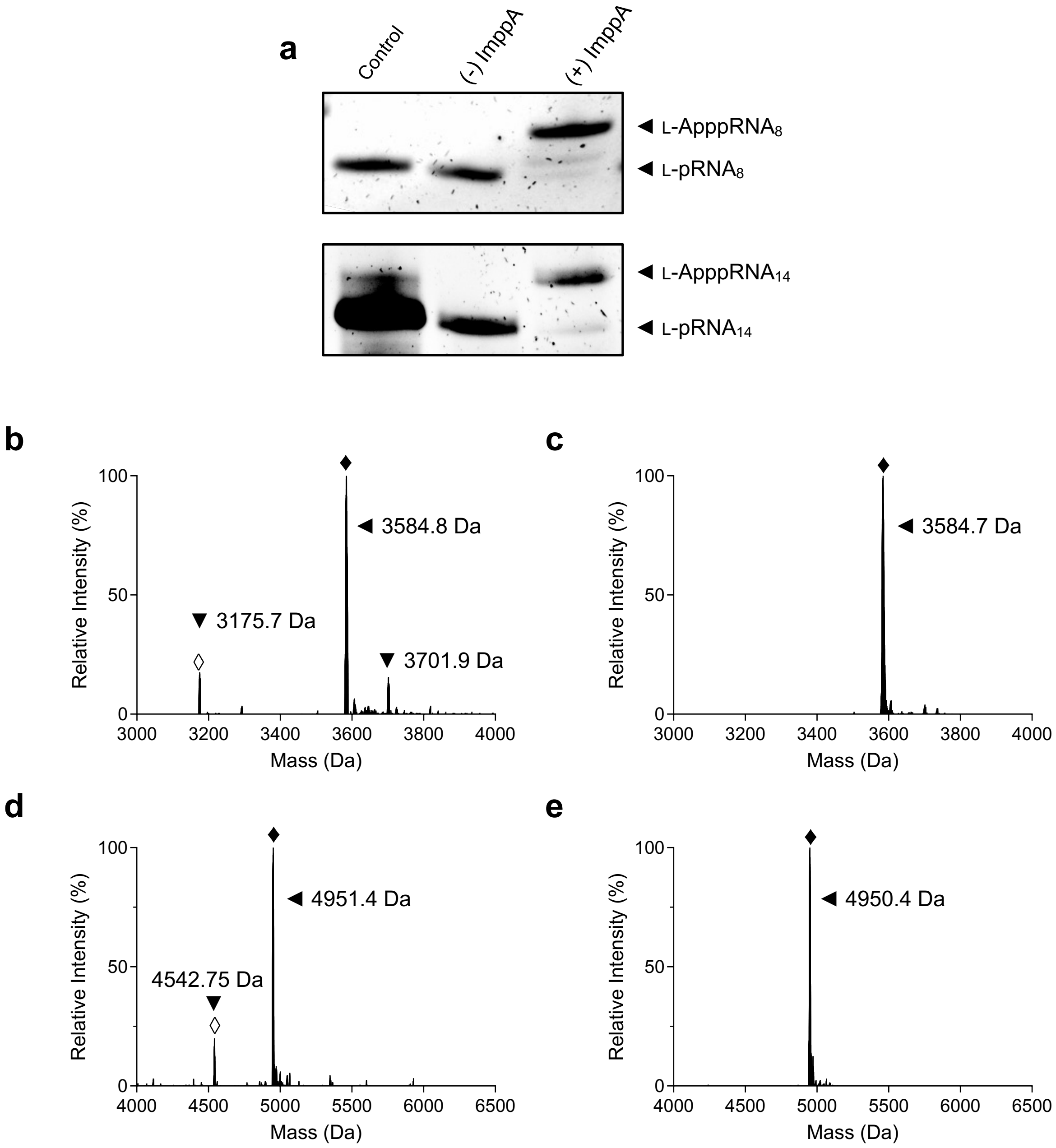


**Figure S5.** (a) Representative denaturing PAGE analysis of the reaction of ImppA with L-pRNA$_8$ (top) and L-pRNA$_{14}$ (bottom). (b) ESI-MS of crude L-ApppRNA$_8$. Calculated: 3584.7 Da, observed: 3584.8 Da. (c) ESI-MS of PAGE purified L-ApppRNA$_8$. Calculated: 3584.7 Da, observed: 3584.7 Da. (d) ESI-MS of crude L-ApppRNA$_{14}$. Calculated: 4950.7 Da, observed: 4951.4 Da. (e) ESI-MS of PAGE purified L-ApppRNA$_{14}$. Calculated: 4950.7 Da, observed: 4950.4 Da. Open diamond = unreacted 5′-phosphorylated starting material; Closed diamond = 5′-adenosyl triphosphate product.

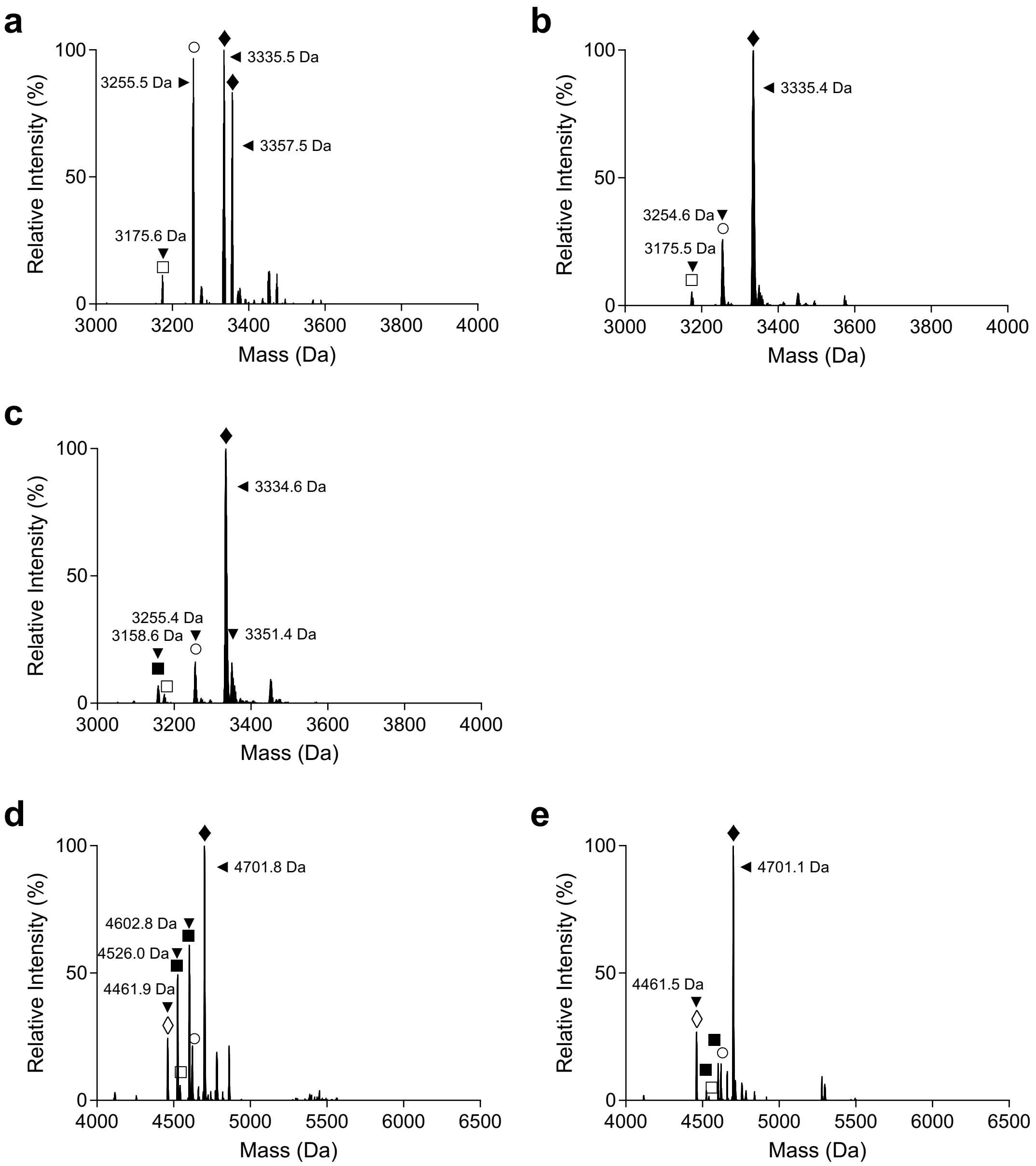


**Figure S6.** ESI-MS characterization of 5′-triphosphorylated L-RNA donors prepared by Ludwig-Eckstein chemistry. (a) ESI-MS of crude L-pppRNA$_8$. Calculated: 3335.2 Da, observed: 3335.5 Da (plus sodium adducts). (b) ESI-MS of PAGE purified L-pppRNA$_8$. Calculated: 3335.2 Da, observed: 3335.4 Da. (c) ESI-MS of HPLC purified L-pppRNA$_8$-CG. Calculated: 3335.2 Da, observed: 3334.6 Da. (d) ESI-MS of crude L-ppp-RNA$_{14}$. Calculated: 4701.1 Da, observed: 4701.8 Da. (e) ESI-MS of PAGE purified L-pppRNA$_{14}$. Calculated: 4701.1 Da, observed: 4701.1 Da. Open diamond = unreacted starting material; Closed diamond = 5′-triphosphorylated product; open square = 5′-monophosphate; closed square = 5′-H-phosphonate; open circle = 5′-diphosphate.

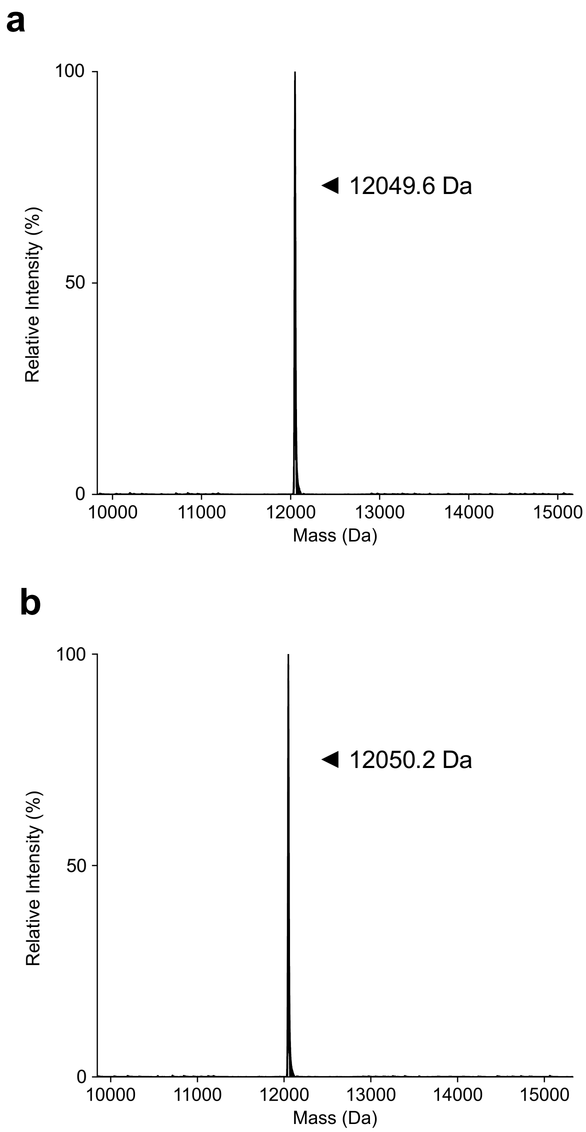


**Figure S7.** (a) ESI-MS data for the ligated product of L-Acceptor$_{hp}$ and L-pppRNA$_8$. Calculated: 12,048.7 Da, observed: 12,048.6 Da. (b) ESI-MS data for the ligated product of L-Acceptor$_{hp}$ and L-ApppRNA$_8$. Calculated: 12,048.7 Da, observed: 12050.2 Da.

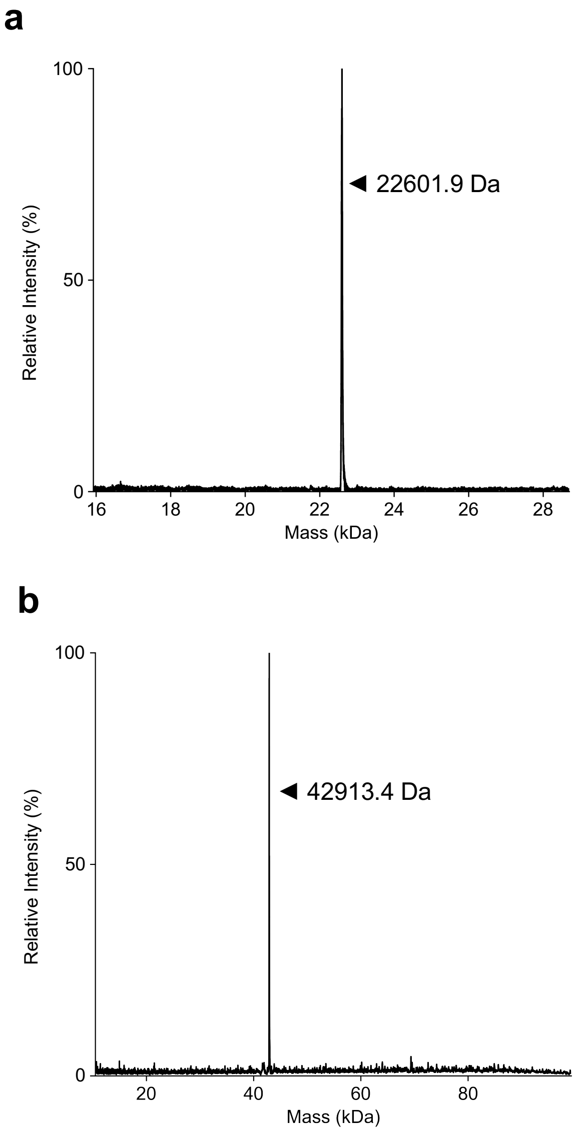


**Figure S8.** (a) ESI-MS characterization of donor L-Appp27.6t$_D$. Calculated: 22598.4 Da, observed: 22601.9 Da. (b) ESI-MS characterization of L-27.6t prepared by cross-chiral ligation. Calculated: 42912.8 Da, observed: 42913.4 Da.

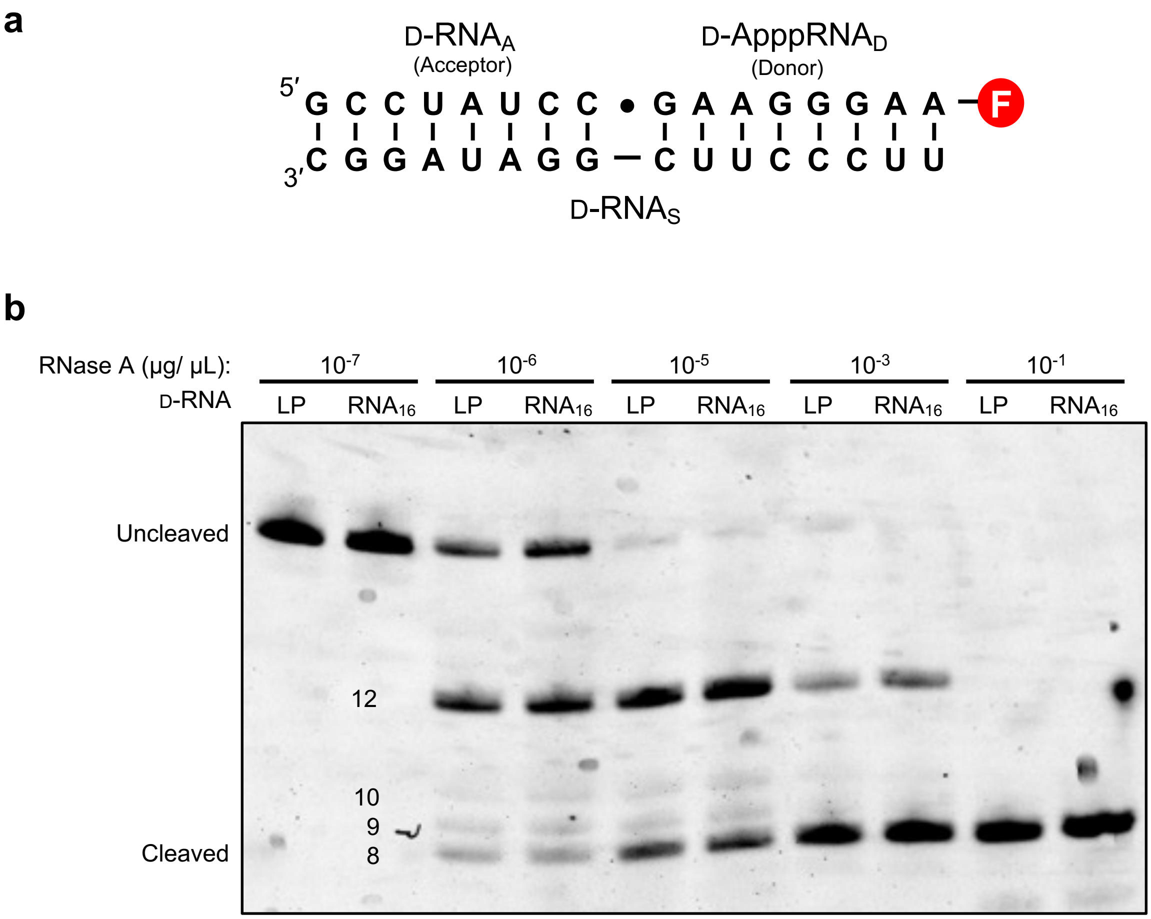


**Figure S9.** Analysis of the regiospecificity of cross-chiral ligation. (a) Sequence and structure of the ligation complex use to assess the regiospecificity of L-27.6t. The ligation junction is indicated by a dot, which is also the RNaseA cleavage site closest to the 3′-end of the product. Circled "F" indicates the dye label (Cy3). (b) Representative denaturing PAGE analysis of the RNase A digestion of the ligated product (LP; D-RNA$_{LP}$) depicted in panel a and the authentic product of the same sequence (D-RNA$_{16}$) demonstrating similar digestion patterns.

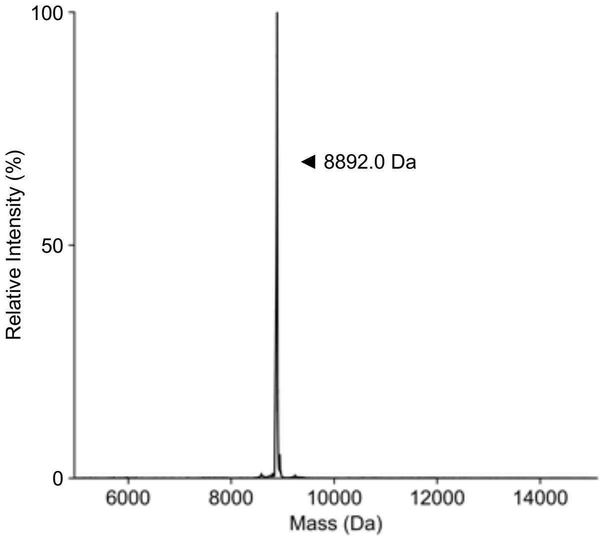


**Figure S10.** ESI-MS characterization of L-Acceptor$_{hp}$ used to generate the ligation complex depicted in Figure 1a. Calculated: 8891.6 Da, observed: 8892.0 Da.

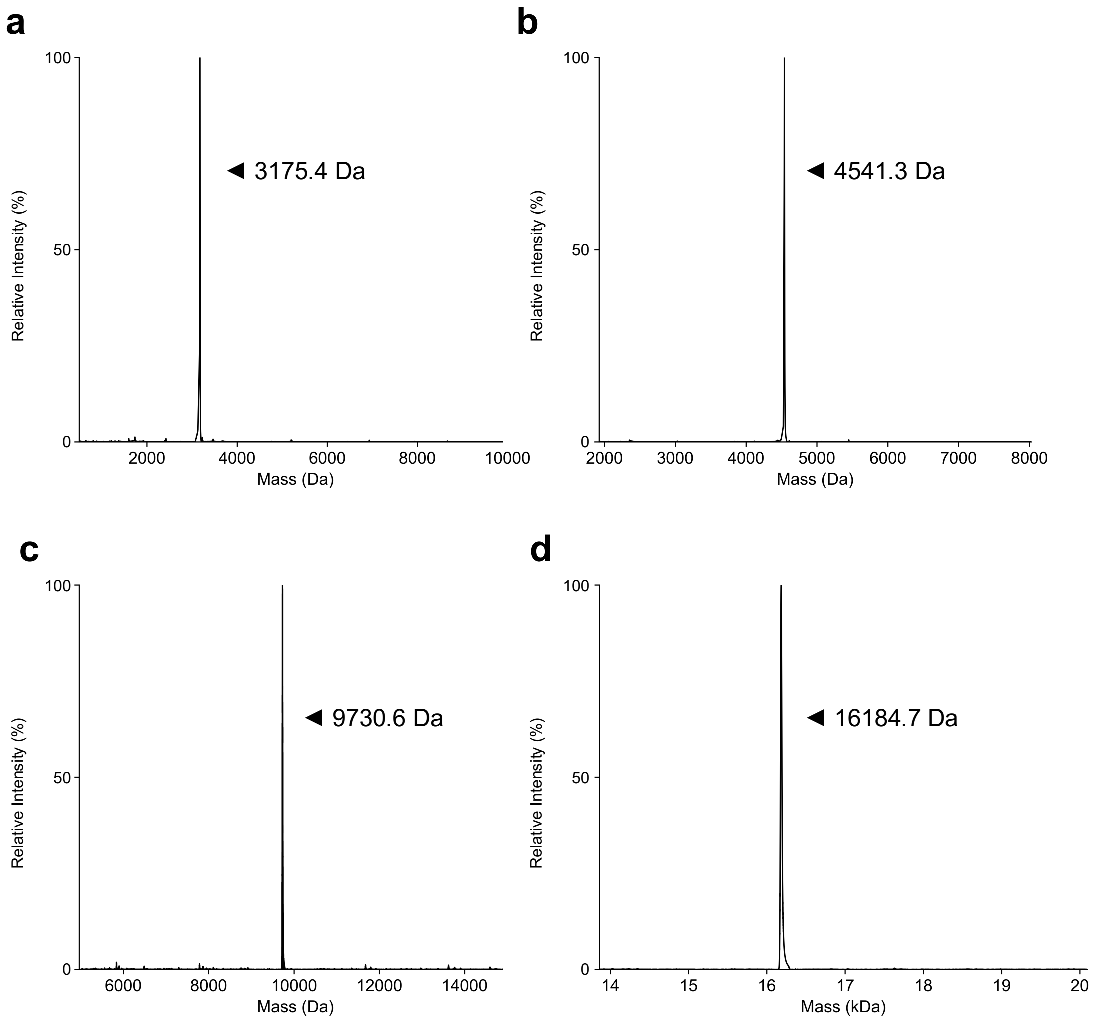


**Figure S11.** ESI-MS of 5′-phosphorylated L-RNAs used to generate donor substrates. (a) ESI-MS of L-pRNA$_8$. Calculated: 3175.1 Da, observed: 3175.4 Da. (b) ESI-MS of L-pRNA$_{14}$. Calculated: 4541.6 Da, observed: 4541.3 Da. (c) ESI-MS of L-pRNA$_{30}$. Calculated: 9730.8 Da, observed: 9730.6 Da. (d) ESI-MS of L-pRNA$_{50}$. Calculated: 16184.6 Da, observed: 16184.7 Da.

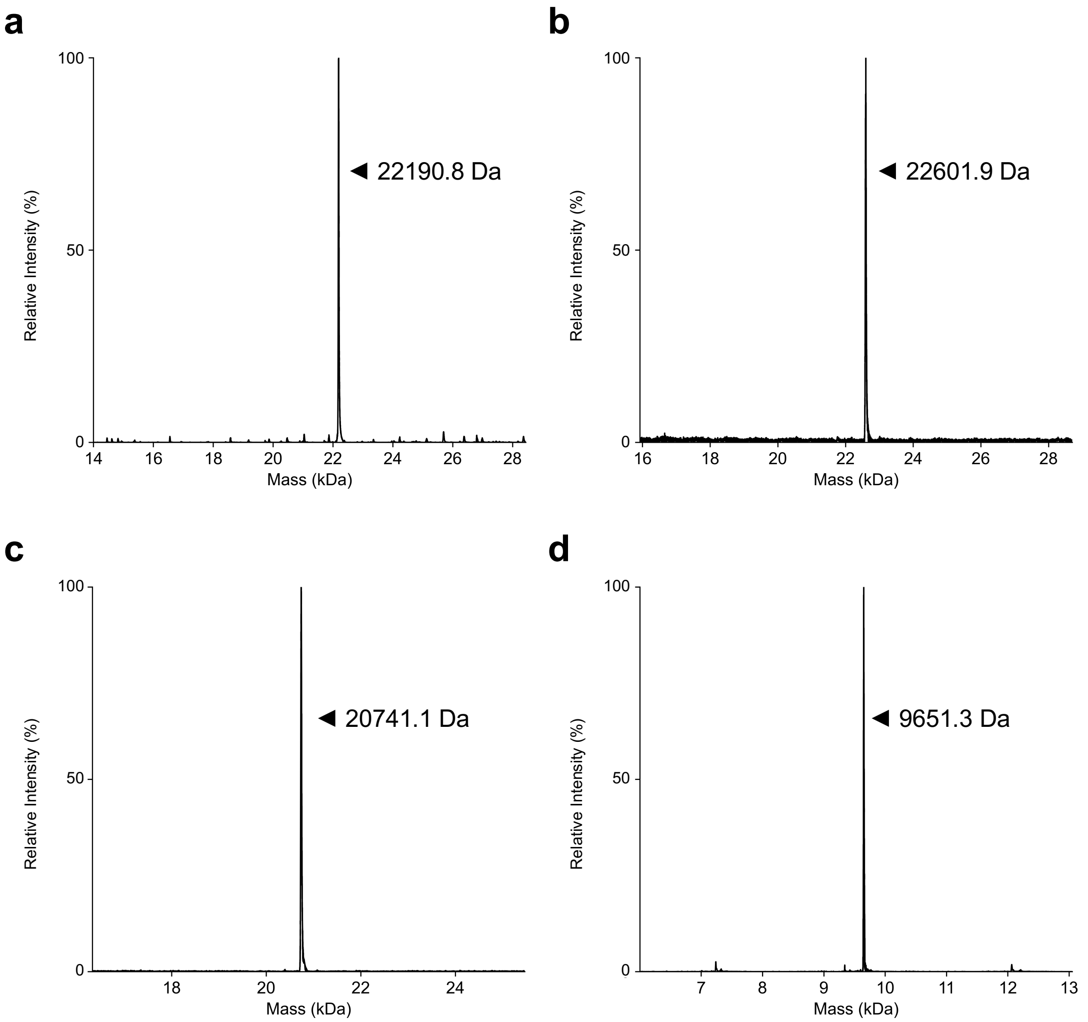


**Figure S12.** ESI-MS for L-RNA substrates associated with the synthesis of L-27.6t. (a) ESI-MS of L-p27.6t$_D$. Calculated: 22189.25 Da, observed: 22190.8 Da. (b) ESI-MS of crude L-Appp27.6t$_D$. Calculated: 22598.4 Da, observed: 22601.9 Da. (c) ESI-MS of L-27.6t$_A$. Calculated: 20741.4 Da, observed: 20741.1 Da. (d) ESI-MS of splint L-27.6t$_S$. Calculated: 9651.7 Da, observed: 9651.3 Da.

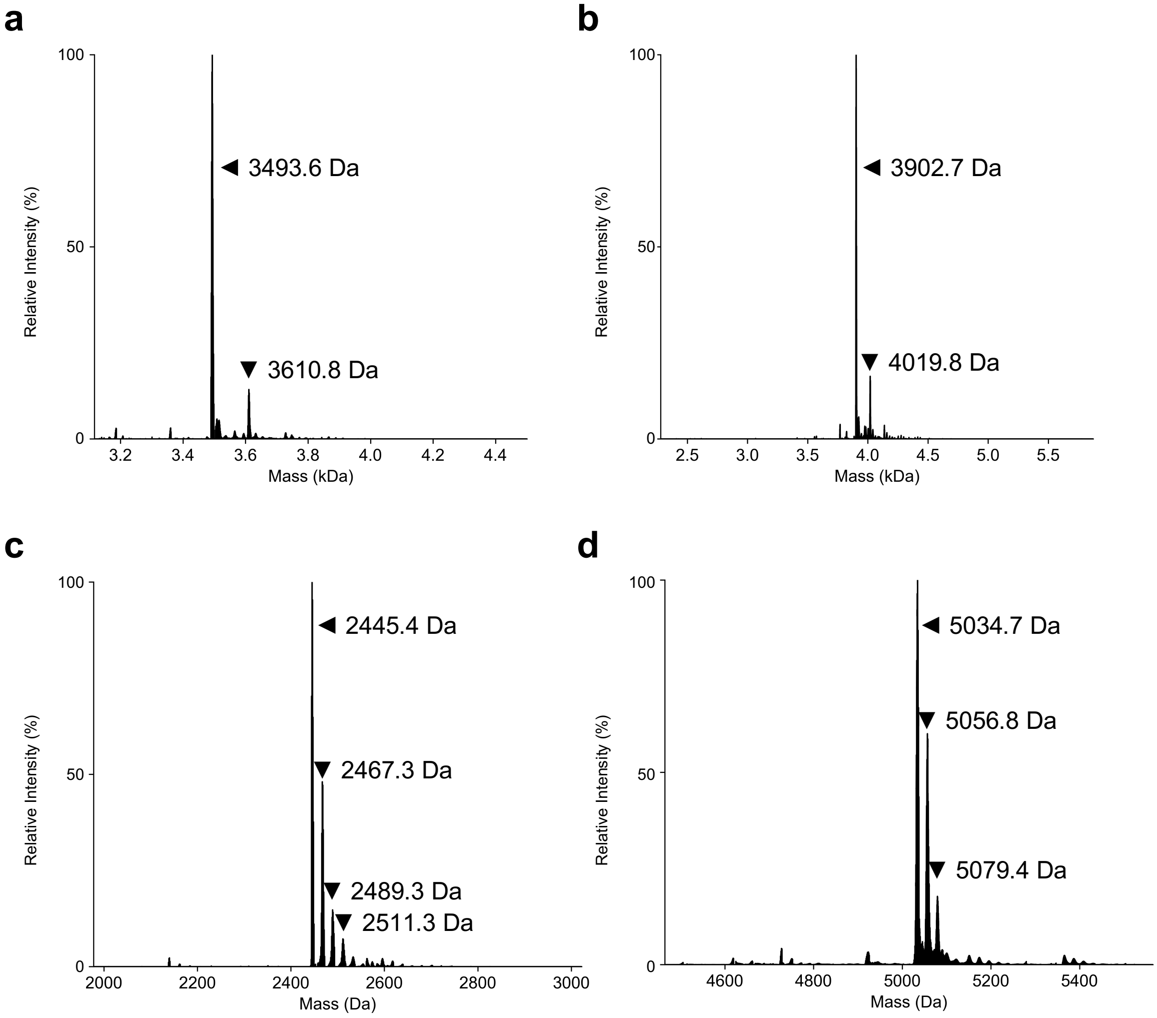


**Figure S13.** ESI-MS characterization of D-RNA ligation substrates depicted in Figure S9a. (a) ESI-MS of D-pRNA$_D$. Calculated: 3492.9 Da, observed: 3493.6 Da. (b) ESI-MS of D-ApppRNA$_D$. Calculated: 3902.0 Da, observed: 3902.7 Da. (c) ESI-MS of D-RNA$_A$. Calculated: 2445.6 Da, observed: 2445.4 Da (plus sodium adducts). (d) ESI-MS of D-RNA$_s$. Calculated: 5034.0 Da, observed: 5034.7 Da (plus sodium adducts).

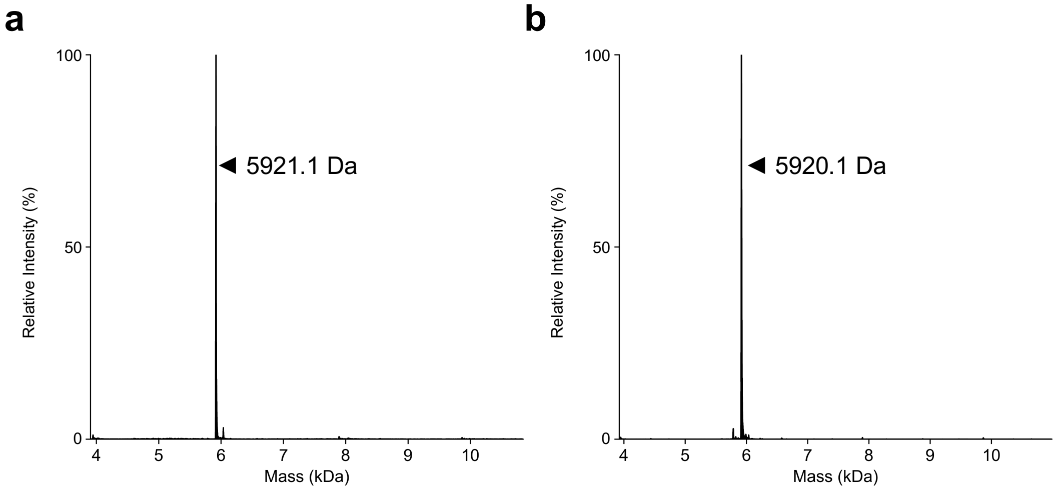


**Figure S14.** ESI-MS characterization of the D-RNA ligation product (LP) and synthetic D-$RNA_{16}$ used in the RNase A digestion analysis (Figure S9). (a) ESI-MS of D-$RNA_{LP}$. Calculated: 5920.9 Da, observed: 5921.1 Da. (b) ESI-MS of D-$RNA_{16}$. Calculated: 5920.9 Da, observed: 5920.1 Da.

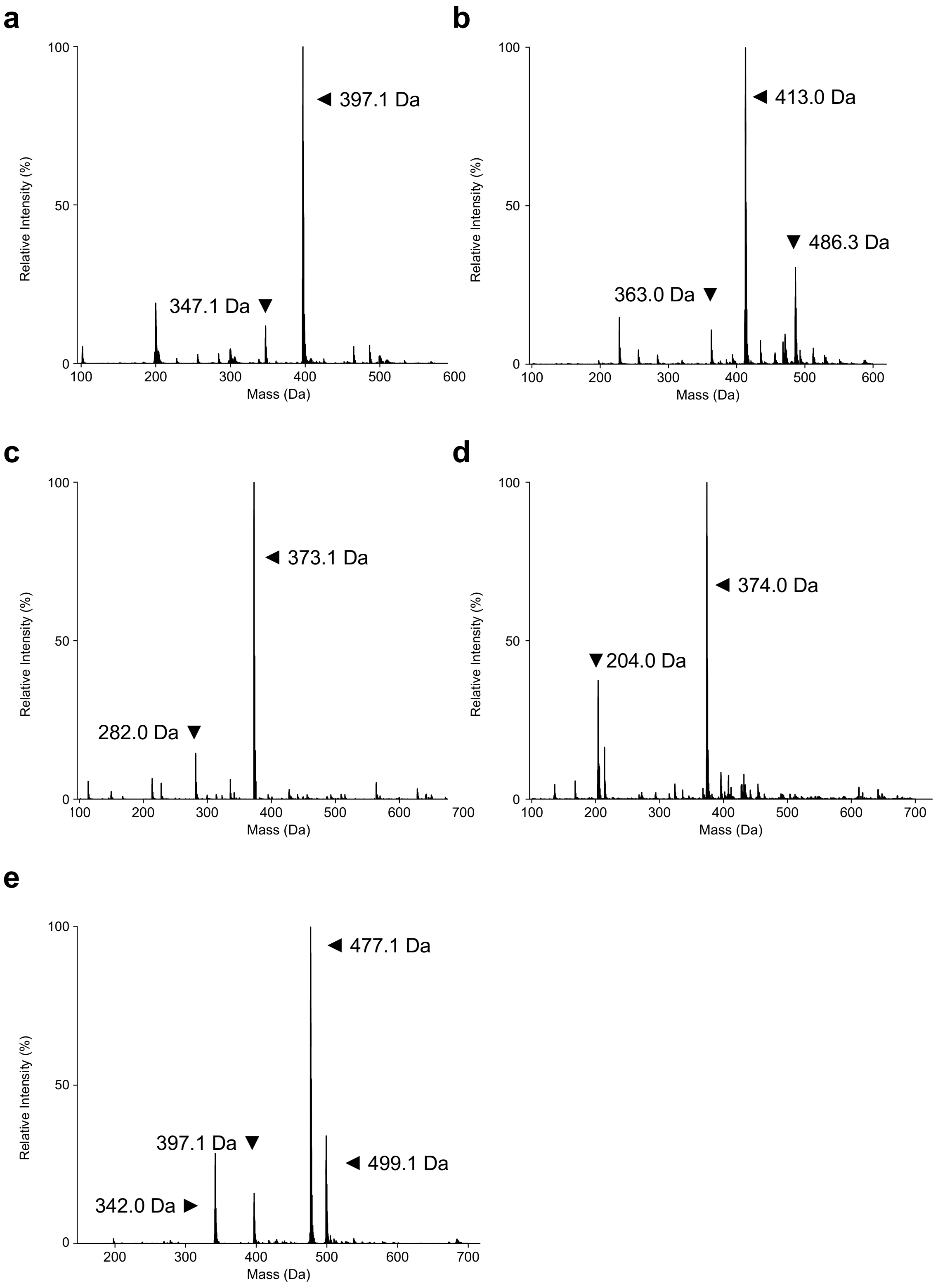

a
397.1 Da
347.1 Da
b
413.0 Da
486.3 Da
363.0 Da
c
373.1 Da
282.0 Da
d
374.0 Da
204.0 Da
e
477.1 Da
397.1 Da
499.1 Da
342.0 Da
Relative Intensity (%)
Mass (Da)

**Figure S15 (previous page).** ESI-MS characterization of crude phosphoimidazolides used in the preparation of 5′-N-ylated RNAs. (a) ESI-MS of ImpA. Calculated: 397.1 Da, observed: 397.1 Da. Estimated purity: 77%. (b) ESI-MS of ImpG. Calculated: 413.1 Da, observed: 413.0 Da. Estimated purity: 57%. (c) ESI-MS of ImpC. Calculated: 373.1 Da, observed: 373.1 Da. Estimated purity: 70%. (d) ESI-MS of ImpU. Calculated: 374.1 Da, observed: 374.0 Da. Estimated purity: 65%. (e) ESI-MS of ImppA. Calculated: 477.1 Da, observed: 477.1 Da (plus sodium adduct). Estimated purity: 75%.

### S3. Supplementary Tables.

**Table S1.** Names and sequences of oligonucleotides used in this work. L-RNA (blue) and D-RNA (black) are indicated by color. /Phos/ = monophosphate; /rNpp/ = N-ylation (N = A,G,C, or U); /rAppp/ = adenosyl triphosphate; /ppp/ = triphosphate; /BioTEG/ = triethylene glycol biotin; /sp18/ = hexaethylene glycol; /AmMC6/ = C6 amino modifier; /6-FAM/ = 6-fluorescein; /Cy5/ = Sulfo-Cy5; /Cy3/ = Sulfo-Cy3.

| Sequence Name | Sequence Identity (5'→3') | ESI-MS (Figure) |
|---|---|---|
| L-pRNA$_{8}$ | /Phos/-GACUGGUC-/BioTEG/ | S11a |
| L-pppRNA$_{8}$ | /ppp/-GACUGGUC-/BioTEG/ | S6b, c |
| L-AppRNA$_{8}$ | /rApp/-GACUGGUC-/BioTEG/ | S1a |
| L-GppRNA$_{8}$ | /rGpp/-GACUGGUC-/BioTEG/ | S3a |
| L-CppRNA$_{8}$ | /rCpp/-GACUGGUC-/BioTEG/ | S3b |
| L-UppRNA$_{8}$ | /rUpp/-GACUGGUC-/BioTEG/ | S3c |
| L-ApppRNA$_{8}$ | /rAppp/-GACUGGUC-/BioTEG/ | S5c |
| L-pRNA$_{14}$ | /Phos/-GACUGGUCAGUCGC | S11b |
| L-pppRNA$_{14}$ | /ppp/-GACUGGUCAGUCGC | S6e |
| L-AppRNA$_{14}$ | /rApp/-GACUGGUCAGUCGC | S1b |
| L-ApppRNA$_{14}$ | /rAppp/-GACUGGUCAGUCGC | S5e |
| L-pRNA$_{30}$ | /Phos/-GACUGGUCAGUCGCAGCAGAAUCGCACUGA | S11c |
| L-AppRNA$_{30}$ | /rApp/-GACUGGUCAGUCGCAGCAGAAUCGCACUGA | S1c |
| L-pRNA$_{50}$ | /Phos/-GACUGGUCAGUCGCAGCAGAAUCGCACUGAAGCACGUGU ACGAUGAUAUU | S11d |
| L-AppRNA$_{50}$ | /rApp/-GACUGGUCAGUCGCAGCAGAAUCGCACUGAAGCACGUGU ACGAUGAUAUU | S1d |
| L-Accpetor$_{hp}$ | /6-FAM/-GACCAGUCGGAUAGCGCAAGCUAUCC | S10 |
| L-p27.6t$_{D}$ | /Phos/-GAUAAAAUGCACAUAGGUCGAAAGACCUUAUACAAGAACU GUAUCACCGGAGGGCGAGCACCACC-/sp18/-/AmMC6/-/Cy5/ | S12a |
| L-Appp27.6t$_{D}$ | /rAppp/-GAUAAAAUGCACAUAGGUCGAAAGACCUUAUACAAGAAC UGUAUCACCGGAGGGCGAGCACCACC-/sp18/-/AmMC6/-/Cy5/ | S12b |
| L-27.6t$_{A}$ | GGUGGCGGACGUGUUUCACGUAUAAUCGUGCGGGACACUGACU CGUCAGUGCAUUGAGAAGGAG | S12c |
| L-27.6t$_{S}$ | CCUAUGUGCAUUUUAUCCUCCUUCUCAAUGC | S12d |
| D-pRNA$_{D}$ | /Phos/-GAAGGGAA-/AmMC6/-/Cy3/ | S13a |
| D-ApppRNA$_{D}$ | /rAppp/-GAAGGGAA-/AmMC6/-/Cy3/ | S13b |
| D-RNA$_{A}$ | GCCUAUCC | S13c |
| D-RNA$_{S}$ | UUCCCUUCGGAUAGGC | S13d |
| D-RNA$_{16}$ | GCCUAUCCGAAGGGAA-/AmMC6/-/Cy3/ | S14b |
| D-16.12t | GGUCGUCAGUGCAUUGAGAAGGAGGAUAAAAUGCACAUAGGUC GAAAGACCUUAUACAAGAACUGUAUCACCGGAGGGCGACC | N/A |
| D-27.3t | GGUGGUGGACGUGAUCAUUACGGAUCACUAACUCGUCAGUGCA UUGAGAAGGAGAAUAAAAUGCACAUAGGUCGAAAGACCUUAUAC AAGAACUGUAUCACCGGAGGGCGAGCACCACC | N/A |
| D-27.6t | GGUGGCGGACGUGUUUCACGUAUAAUCGUGCGGGACACUGACU CGUCAGUGCAUUGAGAAGGAGGAUAAAAUGCACAUAGGUCGAA AGACCUUAUACAAGAACUGUAUCACCGGAGGGCGAGCACCACC | N/A |

**Table S2.** DNA oligonucleotides used in cross extension reactions to generate the dsDNA templates for *in vitro* transcription of the indicated D-ribozymes. The T7 promoter sequence is underlined.

| Ribozyme | Strand | Sequence (5'→3') |
|---|---|---|
| 16.12t | Forward | TTCTAATACGACTCACTATAGGTCGTCAGTGCATTGAGAAGGAGGATAAAATGCACATAGGT |
| | Reverse | GGTCGCCCTCCGGTGATACAGTTCTTGTATAAGGTCTTTCGACCTATGTGCATTTTATCCTCC |
| 27.3t | Forward | TTCTAATACGACTCACTATAGGTGGTGGACGTGATCATTACGGATCACTAACTCGTCAGTGCATTGAGAAGGAGAATAAAATGCAC |
| | Reverse | GGTGGTGCTCGCCCTCCGGTGATACAGTTCTTGTATAAGGTCTTTCGACCTATGTGCATTTTATTCTCCTTCTCAATGCACTGACG |
| 27.6t | Forward | TTCTAATACGACTCACTATAGGTGGCGGACGTGTTTCACGTATAATCGTGCGGGACACTGACTCGTCAGTGCATTGAGAAGGAGGATAAA |
| | Reverse | GGTGGTGCTCGCCCTCCGGTGATACAGTTCTTGTATAAGGTCTTTCGACCTATGTGCATTTTATCCTCCTTCTCAATGCACTGACGAGTC |
